# A Molecular Tension Sensor for N-Cadherin Reveals Distinct Forms of Mechanosensitive Adhesion Assembly in Adherens and Synaptic Junctions

**DOI:** 10.1101/552802

**Authors:** Ishaan Puranam, Aarti Urs, Brenna Kirk, Karen A. Newell-Litwa, Brenton Hoffman

## Abstract

N-cadherin mediates physical linkages in a variety of force-generating and load-bearing tissues. To enable visualization and quantification of mechanical loads experienced by N-Cadherin, we developed a genetically-encoded FRET-based tension sensor for this protein. We observe that N-Cadherin supports non-muscle myosin II (NMII) activity-dependent loads within the adherens junctions (AJs) of VSMCs and the synaptic junctions (SJs) of neurons. To probe the relationship between mechanical loads and AJ/SJ formation, we evaluated the relationships between N-cadherin tension and the size of these adhesion structures. In VSMCs, no relationship between N-cadherin tension and AJ size was observed, consistent with previously observed homeostatic regulation of mechanical loading. In neurons, a strong correlation between SJ size and N-cadherin load was observed, demonstrating an absence of homeostatic regulation. Treatment with glycine, a known initiator of synapse maturation, lead to increased SJ size and N-cadherin load, suggesting a role for mechanosensitive signaling in this process. Correspondingly, we observe that NMII activity is required for the Src-mediated phosphorylation of NMDAR subunit GluN2B at Tyr 1252, which is a key event in synaptic potentiation. Together these data demonstrate N-cadherin tension is subject to cell type specific regulation and that mechanosensitive signaling occurs within SJs.

## Introduction

Within tissues, the primary functions of cell-cell adhesions are to enable structure formation, maintain mechanical integrity, and facilitate signal transmission [1, 2]. These functions require cell-cell adhesions to resist mechanical loads. This is enabled by a variety of sub-cellular structures, such as adherens junctions (AJs) and synaptic junctions (SJs), in different cellular contexts. Traditionally, the mechanical integrity of these structures is thought to be mediated by cadherins, a large family of transmembrane proteins with over 100 members [3]. These proteins simultaneously form calcium-dependent adhesive interactions between cells, and link to the stiff actin cytoskeleton through interactions with catenins and a variety of other scaffolding proteins [1, 4]. However, recent advances in the emerging field of mechanobiology have demonstrated that cell-cell adhesion is not solely mediated by the passive, adhesive interactions of cadherins and other proteins. Instead, tensile forces play a key role in the formation and stabilization of cell-cell adhesions [5-10]. These tensile forces activate mechanosensitive signaling pathways through the alternation of protein conformations and the formation of new protein-protein interactions. Most research has focused on the mechanobiology of AJs formed by epithelial or endothelial cells, which are mediated by the classical cadherins E-cadherin and VE-cadherin, respectively [1, 11]. Excitingly, these studies are revealing new insights into the mechanical aspects of tissue development, wound healing, and the initiation of mechanosensitive diseases including cancer and atherosclerosis; however, the mechanosensitive roles of the other cadherin family members in various tissue types are not as well understood.

In many tissues, cell-cell adhesion is mediated by AJs/SJs containing another classical cadherin, N-cadherin [2, 12, 13]. Loss of N-cadherin results in embryonic lethality due to a wide variety of malformed tissues, including cardiovascular and neural tube defects [14]. Like other cadherins, the primary function N-cadherin-based AJs was thought to be the regulation of adhesion of mesenchymal or neuronal cells through homotypic interactions. Recent evidence has revealed other mechanical functions for these structures. N-cadherin based AJs transmit substantial mechanical loads in a variety of cell types, including cardiomyocytes [15], myogenic cells [16], HGF-treated transformed epithelial cells [17], and within heterotypic cadherin linkages formed between cancer-associated fibroblasts and cancer cells [18]. N-cadherin based AJs also respond to altered mechanical cues by changes in size and adhesion strength [19-21], demonstrating a key role for mechanosensitive process in these structures. Notably, N-cadherin forms homotypic interactions with distinct physical characteristics from other cadherin pairs. Key differences include different affinities and disassembly kinetics in homotypic interactions [22, 23]. These data suggest that N-cadherin functions as a mechanosensitive protein similar to other classical cadherins, but likely has distinct roles due its differential expression pattern and distinct biophysical properties. However, further progress is limited by a lack of tools for probing the mechanics of N-cadherin on the molecular scale. Therefore, in this work we sought to develop, validate, and utilize a novel biosensor for studying the forces experienced by N-cadherin in a variety of cell types.

Specifically, we sought to study mechanical roles of N-cadherin in two systems, one involving homogenous junctions containing similar proteins within the linked membranes of the neighboring cells, and another involving heterogeneous junctions with distinct compositions in the linked membranes. The first system we chose is vascular smooth muscles (VSMCs). These are the most abundant cell in the blood vessel wall and are known to play key roles in a variety of physiological and pathological processes, including blood pressure regulation and neointima formation [24]. Importantly, within the vessel wall, healthy VSMCs physically interact to form a layered, circumferential arrangement that provides structural integrity and changes the diameter of the vessel in response to coordinate changes in VSMC force generation [25, 26]. Remodeling and diseased vessels are commonly characterized by a loss of this organized arrangement and a reduction in the physical linkages between cells, suggesting a key role for VSM cell-cell adhesion in these processes [27, 28]. N-cadherin is the most abundant cadherin in VSMCs and is thought be the primary mediator of mechanical linkages [29]. Consistent with this idea, N-cadherin mediates the formation of homogenous AJs in these cells. These AJs exhibit a large degree of structural variability, ranging from several micron long puncta to 50 micron long ribbon-like structures that run the entire length of the cell depending on the exact cellular context [28, 29]. Furthermore, N-cadherin is implicated in the regulation of VSMC migration, growth, and apoptosis as well as having a key role in the mechanosensitive physiological and pathological process, including the myogenic response, the maintenance of barrier function, and post-injury vasculature repair [28, 30-33]. Thus, a greater understanding of the mechanical loads experienced by N-cadherin in VSMCs is likely to inform several key aspects of cardiovascular physiology and pathophysiology.

For the second system containing heterogenous cell-cell contacts, we chose to focus on the excitatory synapses within rat hippocampal neurons. Excitatory synapses in hippocampal neurons underlie learning and memory formation [34]. Synapses mediate the directional flow of signals from axons to dendrites through the tightly regulated formation of mechanically-coupled, heterogenous pre- and post-synaptic compartments [35]. The majority of excitatory synapses form on specialized post-synaptic dendritic spines that cluster glutamate receptors and adhesion proteins adjacent to the pre-synaptic axon terminal [36]. Changes in excitatory synapse shape and size occur in response to neural activity [37]. For example, during synaptic strengthening, dendritic spines increase the area of the excitatory synapse by adopting a mushroom-shape to increase the membrane surface area adjacent to the pre-synaptic axon terminal [34]. These synaptic changes mediate long-term potentiation (LTP), a persistent strengthening of synapses due to recent neural activity that is a key first step in memory formation [34, 38, 39]. Defective spine morphogenesis and synaptic plasticity in response to neurotransmission are associated with impaired LTP as well as a variety of cognitive defects including neurodevelopmental disorders, psychoses, and neurodegenerative diseases [40].

N-Cadherin is the major cadherin subtype of the brain and is enriched at excitatory synapses [41-43]. Early research into SJs proposed that cadherins stabilize, or “lock-in”, pre- and post-synaptic connections, thereby generating mature synapses [44]. Consistent with this idea, N-cadherin has been demonstrated to mediate mechanical linkages between pre- and post-synaptic membranes, and, through interactions with catenins, enable the local coupling of the cytoskeletons of the two neurons as well as the coordinated regulation of actin dynamics [21, 41]. Furthermore, N-cadherin is a key mediator of synaptogenesis, spine plasticity, neural network formation, and LTP [45-49]. Non-muscle myosin, particularly the myosin IIb isoform, has also been identified as key mediator of synapse plasticity and LTP [33, 50, 51]. Strikingly, in response to inhibition of either N-cadherin or myosin, dendritic spines lose the mushroom-shape indicative of mature SJ and revert to a filopodia-like immature phenotype [50, 52, 53], suggesting a mechanosensitve role for N-cadherin at the SJ. Thus, a greater understanding of the mechanical loads experienced N-cadherin in SJs is likely to inform several key aspects of synaptic plasticity, LTP, and potentially memory formation.

To probe the mechanical force experienced by N-cadherin, we generated, validated, and utilized a novel Forster Energy Transfer (FRET)-based biosensor. In VSMCs, the sensor indicates that N-cadherin experiences non-muscle myosin II (NMII)-dependent forces within AJs but is unloaded in other parts of the cell. Interestingly there is no identifiable relationship between AJ size and N-cadherin tension in this system, likely consistent with previously observed homeostatic regulation of protein load [54, 55]. In neurons, the sensor also reveals the existence of NMII-dependent forces. These forces increase in larger SJ and in response to excitatory NMDA receptor activation, indicating an absence of homeostatic regulation of N-Cadherin load. To address whether synaptic signaling events are altered downstream of NMII-mediated tension, we examined Src family kinase signaling, which has been shown to be activated by mechanical tension in other systems [56, 57]. Using a combination of a SH2-GFP biosensor and phospho-specific antibodies [58, 59], we show that NMII-dependent force generation promotes Src signaling at the synapse, leading to phosphorylation of the NMDA receptor. In total, this work shows that N-cadherin load is subject to cell-specific forms of regulation and that mechanosensitive signaling occurs within synapses, consistent with the emerging idea that the synapse functions as a mechanosensory unit of the brain.

## Results

### Generation and Validation of an N-Cadherin Tension Sensor

Recently, we and others have developed a variety of genetically-encoded FRET-based sensors that report the molecular forces experienced by specific proteins in living cells [8, 55, 60, 61]. These engineered proteins are created by the insertion of a tension sensing module, which is typically comprised of an extensible spring-like domain between two fluorescent proteins (FPs) capable of undergoing FRET, into the protein to be studied (Fig. 1A). As force is applied, the spring-like domain extends, leading to the separation of the FPs and an optically-detectable loss in FRET efficiency. In this work, a tension sensing module containing an extensible domain derived from flagelliform, a component of spider silk, and the mTFP1 and mVenus FRET pair, donor and acceptor FPs, respectively, was used [60]. Notably, the flagelliform-based tension sensing module (TSMod-F) is calibrated, and measurements of FRET efficiency can be converted into the molecular forces experienced by the engineered protein [60].

**Figure 1.**
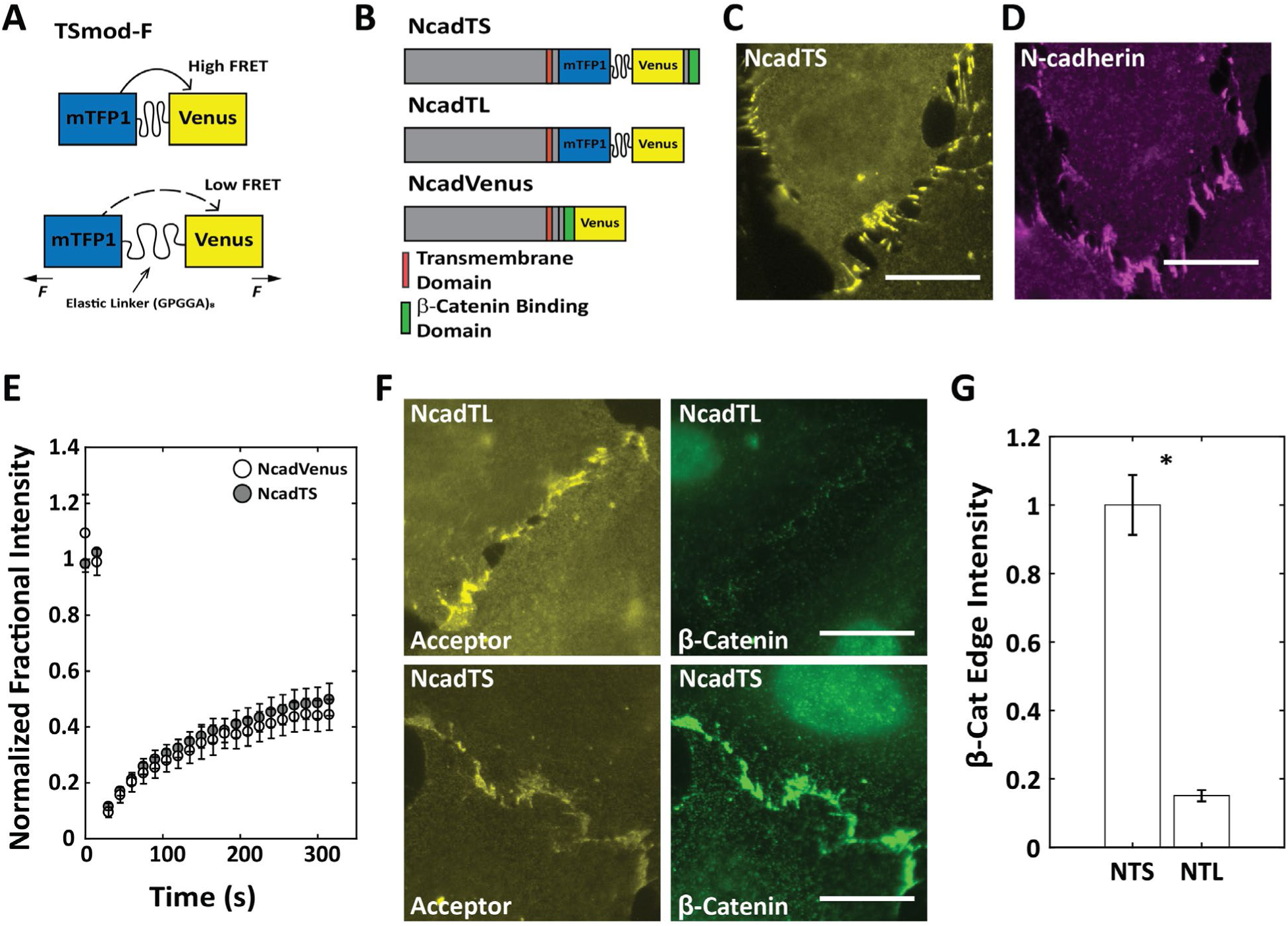
Generation and biological validation of an N-Cadherin Tension Sensor: **A.** The tension sensor module (TSmod-F) consists of two fluorophores separated by an extensible flagelliform linker. When force is applied across TSmod-F, the fluorophores separate, extending the elastic linker and decreasing FRET efficiency. **B.** Schematic of the N-Cadherin Tension Sensor, NcadTS, force insensitive control, NcadTL, and the biological control sensor, NcadVenus. (**C.**-**.D**) Endogenous N-cadherin staining in MOVAS cells mimics localization of transfected NcadTS indicating proper sensor functionality. **E.** FRAP recovery of NCadTS and NCadVenus show no significant differences in the recovery time of the two modules, demonstrating that the insertion of TSmod-F does not disrupt the function of endogenous N-cadherin (n = 24 AJs per group). **F.** β-catenin localization is restored in A10 cells stably expressing NcadTS, but not in A10 cells stably expressing NcadTL, as NcadTL lacks the β-catenin binding site. This indicates that NcadTS is functionally similar to endogenous N-cadherin. **G.** Quantification of β-catenin restoration with NcadTS (NTS) is significantly higher than seen with NcadTL (NTL) (n = 39 and 47 AJs, respectively) (*p < 0.05).

To create an N-cadherin tension sensor (NCadTS), we followed the design of the existing VE-cadherin tension sensor [9]. Specifically, we inserted TSMod-F at amino acid 826 (Fig. 1B). The insertion cite is between the biding sites of p120-catenin, which plays a key role in maintaining cadherins at the membrane, and β-catenin, which in combination with α-catenin mediates linkages to the force-generating actomyosin cytoskeleton [1]. To enable validation of the new biosensor, a variety of control constructs were also developed. A tension-insensitive control senor without the “tail”, or C-terminal region of N-cadherin, was created and termed NCadTL (Fig. 1B). This construct lacks the β-catenin binding site required for α-catenin-mediated linkages to the actomyosin cytoskeleton and is not expected to bear substantial load, as has been observed in control constructs for previously developed tension sensors for various cadherins [8, 9]. To enable probing of N-cadherin dynamics, a C-terminally tagged version of N-Cadherin, termed NCadVenus was also constructed (Fig. 1B).

Next, we determined if NCadTS exhibits behaviors similar to unaltered N-cadherin. First, we compared the localization of NCadTS to endogenous N-cadherin in MOVAS cells, a VSMC line that have previously been used to study N-cadherin mediated cell-cell adhesions and expresses endogenous N-Cadherin [62]. MOVAS cells were either transiently transfected with NCadTS or not transfected and subjected to immunofluorescent labeling of N-Cadherin. The localization of NCadTS (Fig. 1C), as indicated by the acceptor (mVenus) channel, which is independent of FRET and directly proportional to the local concentration, closely mimics the localization of endogenous N-Cadherin in untransfected cells as determined by indirect immunofluorescent labelling (Fig. 1D). For subsequent experiments, we used A10 cells, a rat VSMC line which lacks endogenous N-cadherin expression [62]. Fluorescence Recovery After Photobleaching (FRAP) was used to determine if insertion of TSMod-F affects the dynamics of N-cadherin through analysis of A10 cells that were transiently transfected with either NCadTS or NCadVenus. Recovery dynamics showed no significant differences between NCadTS and NCadVenus (Fig. 1E), consistent with no major perturbations in N-cadherin function due to the incorporation of TSMod-F.

To increase the throughput of subsequent experiments, A10 cells were stably reconstituted with either NCadTS or NCadTL using lentiviral transduction and subjected to FACS analysis to identify cells with similar expression levels. Western blot analysis shows that both proteins were produced as a single band at the expected molecular weight (Sup Fig 1A). The ability of NCadTS to mediate AJ formation in cells lacking N-cadherin was probed in A10 cells through the ability to recruit β-catenin, as was done to validate a previously designed tension sensor for E-cadherin [8]. Notably, AJ restoration does not occur in cells expressing NCadTL, which lacks the β-catenin binding site, but does occur in cells expressing NCadTS (Fig. 1F,G). Previous work has shown that a vinculin tension sensor is robust to fixation [63]. As fixation eases requirements for cell imaging and enables the use of blebbistatin to probe the role of myosin-dependent force, we sought to determine if NCadTS is compatible with standard fixation protocols. To do so, NCadTS and NCadTL were evaluated in either live A10 cells or cells that had been fixed in 4% PFA. No significant difference is seen as a result of PFA fixation for either NcadTS or NcadTL (Sup Fig. 1B,C).

For accurate interpretation of the tension sensors, FRET must occur intramolecularly and not intermolecularly. To enable detection of intermolecular FRET, A10 cells were stably reconstituted with control constructs harboring a point mutation that renders either mTFP1 or mVenus non-fluorescent, or “dark” [9]. There constructs are termed NCadTS-dmTFP1 and NCadTS-dVenus, respectively (Sup Fig. 2A). Successfully transduced cells were isolated through FACS and cells were fixed and subjected to FRET imaging. Cells with expression levels approximately equivalent to the those of the analyzed NCadTS expressing cells and having an approximately 1:1 expression ratio of the two constructs were selected through analysis of the mVenus intensity and the ratio of donor to acceptor concentration [64]. FRET efficiencies of less than 5% are observed using the intermolecular FRET controls (Sup. Fig. 2C), which is a commonly assumed experimental noise for these types of experiments [65].

**Figure 2.**
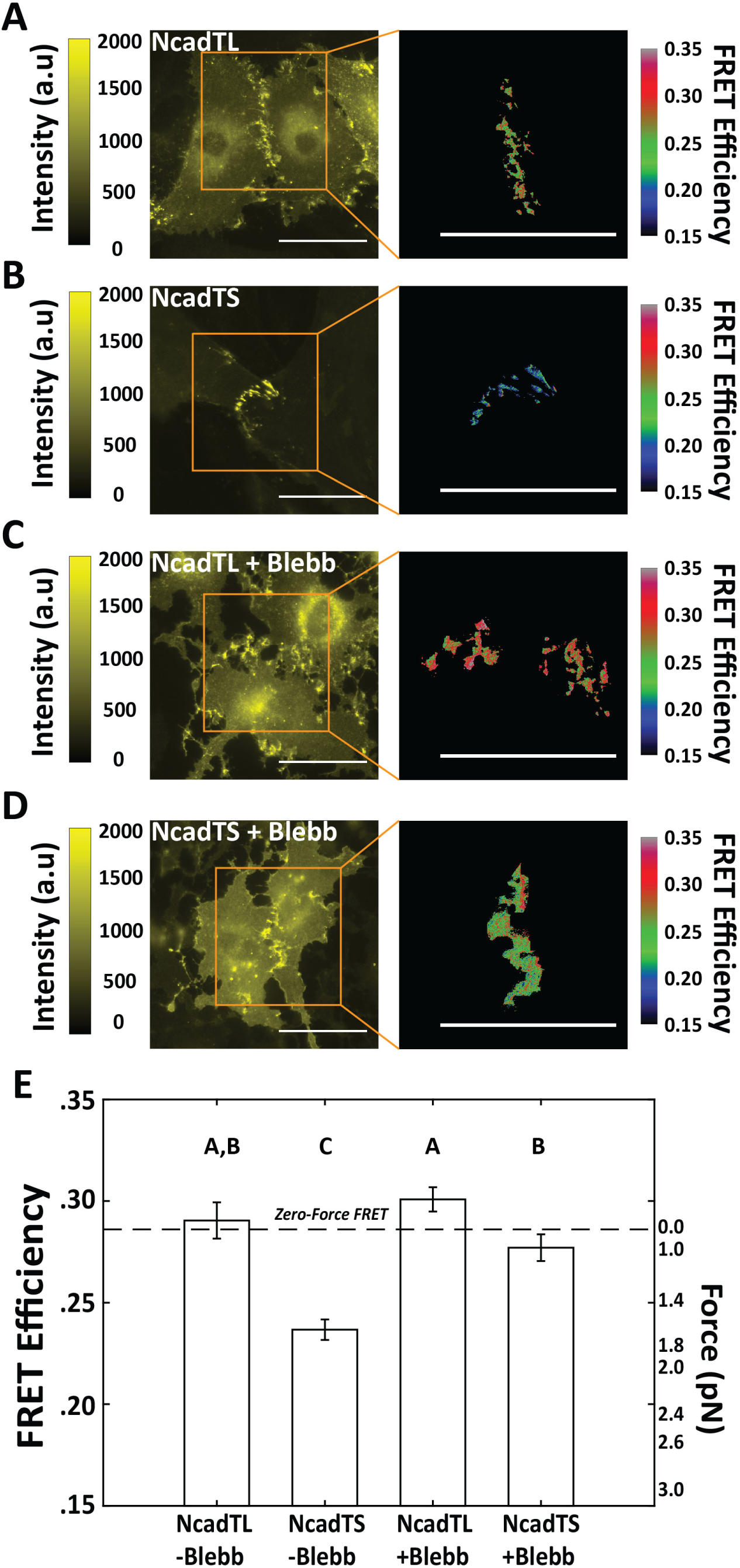
Technical validation of an N-Cadherin Tension Sensor. A10 cells were stably transduced with NcadTL and NcadTS and plated on fibronectin covered glass dishes. **A.** A10 cells expressing NcadTL demonstrated FRET efficiency measurements consistent with a lack of mechanical loading (28.6%), as expected. **B.** A10 cells expressing NcadTS demonstrated a statistically significant decrease in FRET efficiency, corresponding with force across the sensor. **C.** Treatment of A10 cells with Blebbistatin (120 min at 25μM), a cell permeable inhibitor of non-muscle myosin II, did not affect the FRET readings of NcadTL. **D.** Blebbistatin treatment of A10 cells expressing NcadTS demonstrated an unloading of force consistent with a loss of mechanical loading. **E.** Quantification of total data (n = 50, 31, 40, and 30 AJs, respectively). Multiple comparisons are run through non-parametric analysis using the Steel-Dwass test. Experimental groups denoted with the same letter are statistically similar, whereas groups with different letters are statistically different. Specific p-values are reported in Sup. Table 1.

In total, these data show that insertion of TSMod-F into N-cadherin does not affect the localization, dynamics, or ability to interact with key binding partners, demonstrating the ability NCadTS to accurately reflect the characteristics of N-cadherin. Furthermore, the sensor can be used in fixed cells and no intermolecular FRET was observed. Thus, measurements of FRET efficiency from NCadTS can be used to determine the molecular forces by N-Cadherin.

### N-Cadherin Experiences Non-Muscle Myosin II-dependent Forces within the AJs of VSMCs

To evaluate response of the sensor to alteration in cellular force generation, confluent A10 cells stably expressing either NCadTS or NCadTL were fixed, subjected to FRET imaging, and the apparent loads within AJs were determined. Expectedly, NCadTL exhibits FRET Efficiencies near 28.6% corresponding with our previously established “zero-force” state [66] (Fig. 2A,E). NCadTS exhibited statistically significantly lower values of FRET Efficiency, 23.6%, consistent with the tension sensor experiencing average loads of 1.65 pN (Fig. 2B,E). Furthermore, NCadTS and NCadTL localized outside of AJs were not subject to significant mechanical loads (Sup Fig. 3A,B). Thus N-cadherin does not appear to be constitutively loaded, as has been observed in E-Cadherin [8]. To determine the cause of the mechanical loading, cells were treated with blebbistatin, a potent inhibitor of NMII ATPase activity. NMII inhibition did not alter FRET in the tension insensitive NCadTL control (Fig. 2C,E), but resulted in the complete unloading of NCadTS (Fig. 2D,E). Previous work in cadherin mechanobiology has investigated the relationship between the forces at junctions and the size of an AJ, with strong correlation being observed in some systems. N-cadherin tension was not correlated with AJ size or the amount of N-cadherin with a junction (Sup Fig 4). In total, these data demonstrate that NCadTS is capable of reporting molecular forces in living cells and show that N-Cadherin within AJs of VSMCs is loaded by NMII-dependent forces.

**Figure 3.**
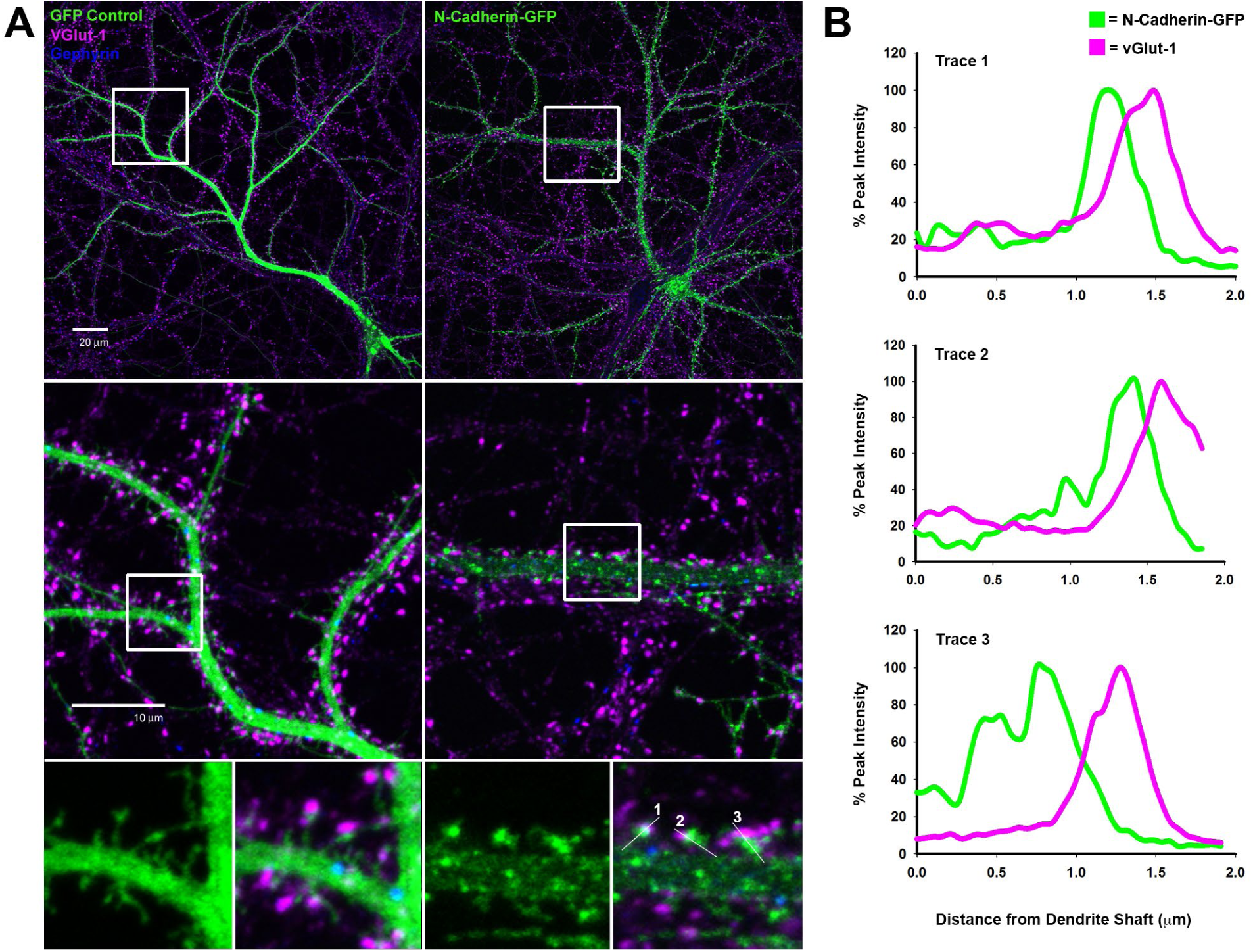
N-Cadherin Localizes to Excitatory Synapses. **A.** Expression of N-Cadherin-GFP (right) in rat primary hippocampal neurons demonstrates that N-Cadherin predominantly localizes to excitatory synapses adjacent to the excitatory pre-synaptic marker, vesicular glutamate transporter-1 (VGlut-1, magenta), and in contrast to the dendritic localization of the inhibitory post-synaptic marker, gephyrin (blue). **B.** Intensity plots of line traces through dendritic spines demonstrate that N-Cadherin is enriched at puncta adjacent to the excitatory pre-synaptic marker, vGlut-1, confirming its localization to excitatory synapses. Line traces correspond to spines identified in the bottom right of panel A.

**Figure 4.**
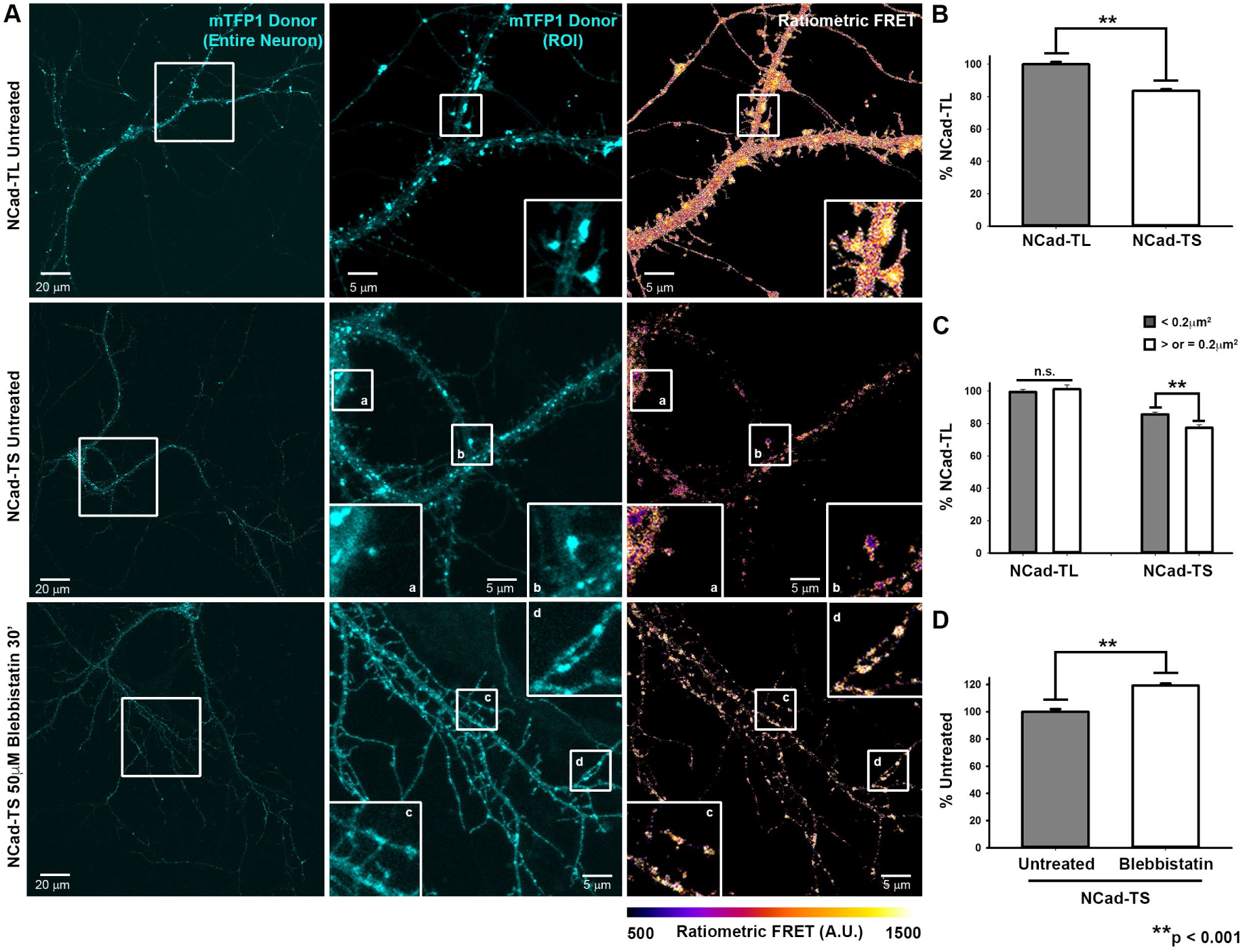
NMII generates forces at excitatory synapses. **A.** NcadTL and NcadTS exhibit punctate staining at spine heads. However, NCadTS, exhibits decreased FRET, indicative of synaptic tension, especially in larger spine heads (panel b). Treatment with 50 μM Blebbistatin for 30 min normalized the FRET signal throughout the neuron, demonstrating that NMII generates synaptic forces. **B.** Quantification of ratiometric FRET in synaptic N-cadherin puncta for untreated NCadTL and NCadTS. (n = 404 and 498 puncta, respectively). All analyzed puncta are ≥ 0.01 mm^2^. Data is normalized to the average FRET for untreated NCadTL. **p<0.001 for untreated NCadTS vs NCadTL, Mann-Whitney Rank Sum Test. **C.** Quantification of ratiometric FRET by puncta size (<0.2mm^2^ or ≥0.2mm^2^). FRET is significantly decreased in larger, more mature (≥0.2mm^2^) NCadTS puncta, indicating an increase in force. NCadTL, which lacks actin association, shows similar FRET regardless of size. (n = 289 NCadTL puncta <0.2mm^2^, 115 NCadTL puncta ≥0.2mm2), n.s., Mann-Whitney Rank Sum Test, and (n = 371 NcadTS <0.2mm^2^ and127 NCadTS puncta ≥0.2mm^2^) **p<0.001, Mann Whitney Rank Sum Test. **D.** Quantification of ratiometric FRET for untreated NCadTS vs Blebbistatin-treated NCadTS (n = 149 and 170 puncta, respectively). Mann-Whitney Rank Sum Test. Data is normalized to the average FRET for untreated NCadTS. **p<0.001, Mann Whitney Rank Sum Test.

### Increased Non-myosin II-dependent N-Cadherin Tension is Associated with Synapse Maturation

To probe the applicability of NCadTS to other systems we sought to determine whether N-cadherin experiences forces within synapses formed by rat primary hippocampal neurons. A C-terminal N-Cadherin-GFP construct predominantly localizes to excitatory glutamatergic synapses, where it is concentrated at the tip of post-synaptic dendritic spines (Fig. 3 A,B) [67]. NCadTS and NCadTL exhibit a similar synaptic localization (Fig. 4A, see mTFP1 donor image), demonstrating the constructs exhibited the expected behavior in this system. To limit the number of required control constructs and simplify analyses of these synaptic puncta, we used ratiometric imaging to quantify FRET. This type of imaging and analysis produces a semi-quantitative FRET Index which, unlike FRET Efficiency measurements, cannot be used to determine the forces experiences by the sensor [68]. However, the indexes can be used to determine relative changes in FRET and the loads experienced by tension sensors within the sensitivity limits of the sensor, which are known to be 1-6 pN for the sensor used in this work. [60].

For NCadTS, the FRET index decreased to ∼83.6% of that observed in NCadTL, demonstrating that N-Cadherin is under force at synapses of rat hippocampal neurons (Fig. 4B). Furthermore, we observed decreased NCadTS FRET primarily in larger, more mature spines (Fig. 4A, compare mushroom-shaped dendritic spine in box b with the less mature dendritic spine in box a). We quantified N-Cadherin puncta size as a readout for spine maturation based on our previous observation that the average post-size of synaptic density is approximately 0.2 μm^2^ in rat hippocampal neurons, but increases with NMII activation [50]. N-Cadherin puncta with sizes greater than or equal to 0.2 μm^2^ have significantly lower FRET than smaller puncta, while no size dependence is observed in synapses containing NCadTL (Fig. 4C). As our previous research demonstrated that NMII drives the shape changes association with spine maturation [50, 53] we therefore hypothesized that NMII generates the observed loading of N-cadherin within synapses. Blebbistatin significantly increased NCadTS FRET (Fig. 4D), identifying NMII as the cause of mechanical loading. Overall, these data demonstrate the NMII-generated forces load N-cadherin with synapses and that these loads are increased in larger, more mature synapses.

### Excitatory Stimulation Increases Force Generation

Excitatory stimulation of NMDA receptors promotes spine maturation and synapse formation, strengthening the synapse during LTP [34]. NMDAR-driven synaptic strengthening also requires NMII activity [50, 53]. We therefore hypothesized that excitatory stimulation of NMDA receptors would increase tension across N-Cadherin, resulting in decreased FRET by NCadTS. Glycine exposure, a process often referred to as chemical LTP, was used to stimulate the NMDA receptor [69], and resulted in mature mushroom-shaped spines with larger N-Cadherin puncta (Fig. 5 A,B). Chemical LTP increased tension across N-Cadherin as demonstrated by a significant decrease in FRET when compared with untreated controls (Fig. 5C), consistent with the increased mechanical loading of N-cadherin in response to neuronal activation.

**Figure 5.**
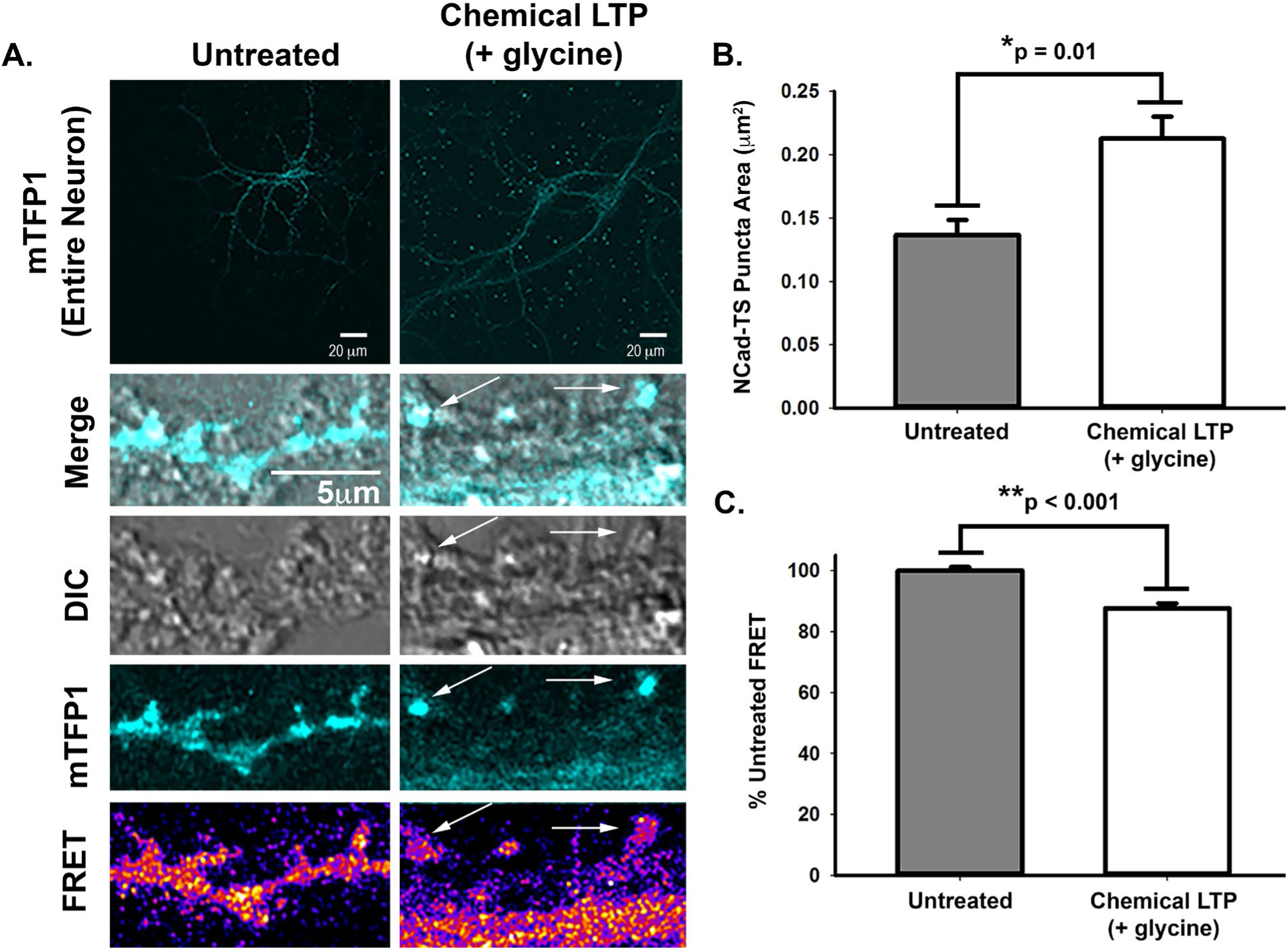
Excitatory stimulation drives force production together with synapse maturation. **A.** Representative images of NCadTS-expressing neurons untreated or treated with glycine to activate NMDA receptors, leading to LTP (Park *et al.*, 2004). Note the large NCadTS puncta that form in response to glycine-mediated excitatory stimulation (arrows). **B.** Quantification of N-Cadherin puncta size for untreated and glycine-treated neurons. Glycine significantly increases N-Cadherin puncta size (n = 268 and 297 puncta, respectively). *p = 0.01, Mann-Whitney Rank Sum Test. **C.** Quantification of NCadTS FRET in untreated vs. glycine-treated neurons. Data is presented as a percent of the average untreated control FRET (n = 268 and 297 puncta, respectively). Glycine significantly decreases FRET. **p<0.001, Mann-Whitney Rank Sum Test.

### Myosin-II-mediated Force Generation Regulates Src Signaling at Synapses

Src-family kinases are known to play a key role in regulation of NMDAR activity during synaptic potentiation[70] and is often involving in mechanosensitive signaling [56]. Therefore, we sought to determine whether NMII activity regulates synaptic Src family kinase signaling. To do so, we used a fluorescent Src SH2-GFP biosensor to visualize the location of active Src in rat hippocampal neurons [58]. Src SH2-GFP localizes to excitatory synapses (Fig. 6A). Y-27632-mediated NMII inhibition reduces this synaptic enrichment, as visualized through time-lapse imaging of the resulting displacement of the SH2-GFP from the synapse coincident with reversion to filopodia-like spine precursors (Movie 1). As the Src-mediated phosphorylation of the NMDAR subunit GluN2B at Tyr 1252 potentiates Ca^2+^ influx important for synaptic potentiation [71], we sought to determine, if this key event is regulated by NMII activity. In untreated samples, NMDAR pTyr1252 localizes to excitatory synapses (Fig. 6B). Blebbistatin treatment reduces GluN2B pTyr1252 puncta size (Fig. 6C), consistent with NMII-mediated regulation of synaptic Src signaling events.

**Figure 6.**
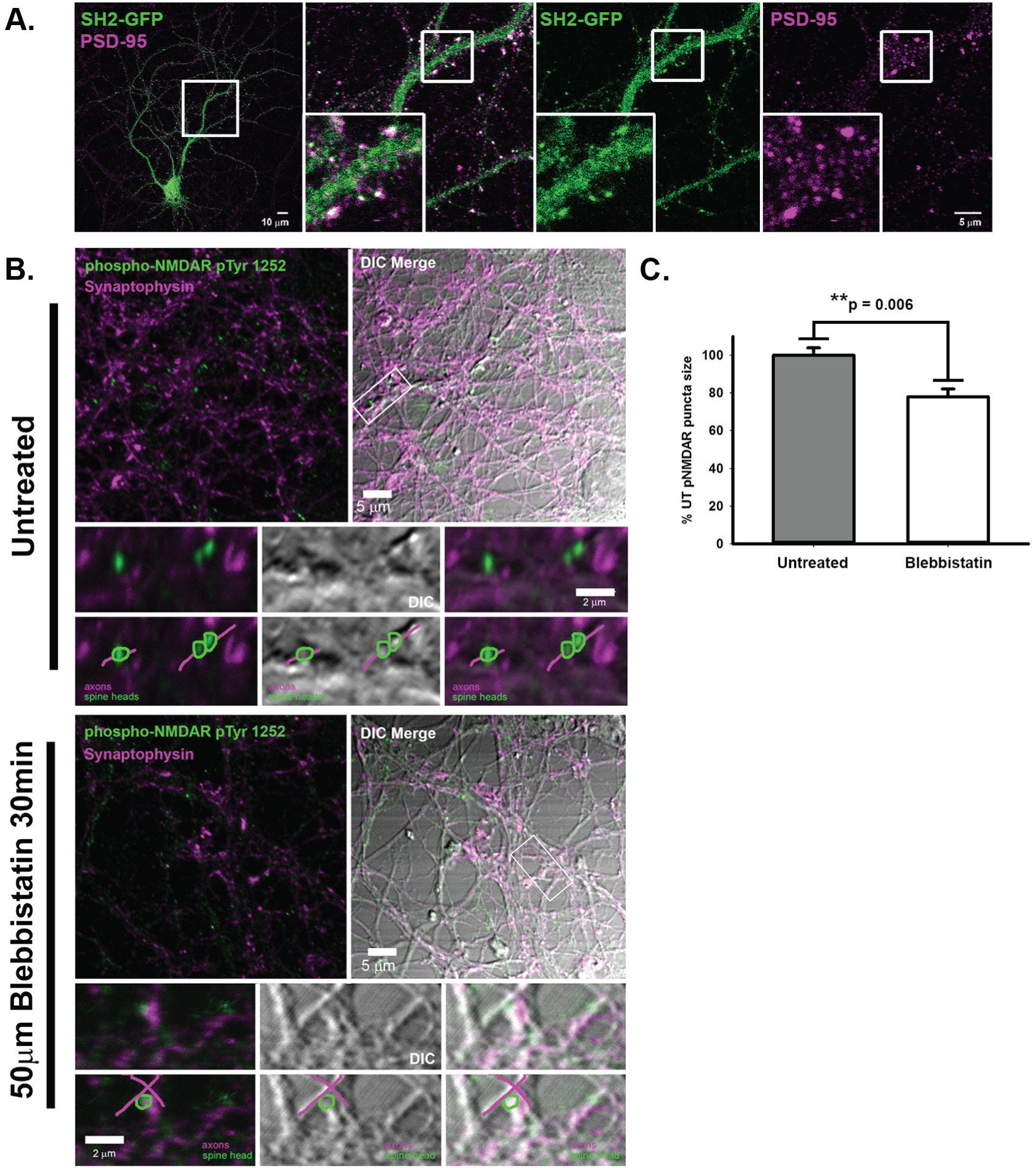
NMII Regulates Src Signaling at the Synapse. **A.** A Src SH2-GFP biosensor localizes to post-synaptic dendritic spines as shown by overlay with post-synaptic density-95 (PSD-95). **B.** Rat hippocampal neurons were treated with 50μM blebbistatin for 30minutes, and stained for the Src post-synaptic substrate, NMDAR pTyr1252, together with the pre-synaptic marker, Synaptophysin. **C.** Blebbistatin treatment reduces NMDAR pTyr1252 post-synaptic area. (n = 1232 and 769 puncta, respectively). **p =0.006, Mann-Whitney Rank Sum Test.

## Discussion

In this study, we developed a genetically encoded FRET-based tension sensor for N-Cadherin, the classical cadherin most commonly found in a variety of load-bearing or force-generating cell types, including fibroblasts, chondrocytes, smooth muscle cells, cardiomyocytes, and neurons. Specifically, we investigated the forces experienced by N-cadherin within AJs in VSMCs, and SJs in neuronal synapses. While the loading of N-cadherin was dependent on NMII-activity in both systems, the relationships between AJ/SJ size and N-cadherin load were distinct. In VSMCs, no relationship between AJ size and N-cadherin load was observed. In neurons, we observe that larger SJs, as well as those that were induced to maturity through glycine exposure, exhibit larger tensions. Furthermore, we demonstrate that NMII activity is required for normal Src signaling in the SJs. Together this data demonstrate that N-cadherin can mediate cell-type specific mechanical responses and that mechanosensitive signaling occurs with SJs.

Typically, mechanical loading of cadherins is thought to require trans-interactions between cadherins on adjacent cells as well as mechanical linkages to the to the force generating actomyosin cytoskeleton [72]. Consistent with this notion, mechanical loading of N-cadherin was limited to areas within cell-cell contacts, where trans-interactions between cadherins and connections of cadherin cytoskeleton are enriched. In contrast, the first measurements of the forces of E-cadherin in MDCK cells with a molecular tension sensor revealed constitutive loading both inside and outside of AJs [8]. Notably, measurements of VE-cadherin loading in endothelial cells revealed loading only within AJs [9], consistent with requirements for trans-interactions and linkages to the actomyosin cytoskeleton. Direct comparison of these data is complicated by a variety of factors, including the fact that the tension sensors have different designs. In the E-cadherin sensor, the TSMod-F is placed between the transmembrane domain and the p120-catenin binding site [8]. In the VE-Cadherin and N-cadherin sensors, the TSMod-F was placed between the p120-catenin and β-catenin binding sites [9]. Recent advances have demonstrated that different localizations within the same protein can support differing loads [73]. Therefore, the presence or absence of constitutive cadherin loading could be due to differences in the various cadherins, cell types, or sensor design. Distinguishing between these possibilities could reveal important aspects of tissue-specific AJ mechanobiology and is an important aspect of future work.

Now pioneering work has clearly established that the E-cadherin and VE-cadherin containing AJs within epithelial and endothelial cells respond to mechanical force [7, 74]. Common responses are changes in assembly dynamics of AJs, often referred to as mechanosensitive assembly or adhesion strengthening, and the activation of mechanosensitive signaling pathways [4]. These mechanosensitive responses can be initiated by externally-applied forces, such fluid shear stress, as well as the internal-generated forces due to actin polymerization or NMII activity. Here we observed that N-cadherin bears load in multiple types of adhesion structures. Previous work has shown that N-cadherin recruitment to SJ is increased during maturation [42]. Here, we showed that N-cadherin loads also increase during maturation. Additionally, acute inhibition of NMII destabilizes synaptic connections [33, 50]. Furthermore, loads increase in response to synapse maturation, and are required for Src-mediated phosphorylation of GluN2B at Tyr 1252, a key event in synaptic potentiation. In combination, these data suggest that mechanosensitive assembly and signaling occur in SJs and may play a key role in physiological processes like LTP.

Mechanosensitive assembly is often understood as a form of homeostatic regulation, where adhesion assembly increases in response to an applied load to maintain a constant stress or force per molecule [54, 75]. The mechanosensitive nature of adhesion structures can be probed through analysis of loads in a structure and the size of the structure. Analysis of molecular loads supported by E-cadherin within AJs of epithelial cells was constant across adhesion of various sizes, consistent with homeostatic regulation [54]. An analysis of the relationship between the size of the AJs in VSMCs and N-cadherin tension revealed no correlation, consistent with the homeostatic models of adhesion strengthening. In neurons we observed strong correlations between the size of a SJ and the mechanical loads experienced by N-cadherin, suggesting a distinct form of regulation. Complete elucidation of the physical bases for this discrepancy will require future work. Likely possibilities include insufficient N-cadherin recruitment to redistribute increased loads or alterations in the cadherin-catenin complex, or other components of the SJ, to affect the way N-Cadherin supports load.

A key next step will be further elucidating the mechanosensitive signaling pathways activated by the loading of AJs and SJs. Notably, the presence or absence of homeostatic mechanisms in adhesion structures would likely affect the dynamics of mechanosensitive signaling. Adhesion structures subject to homeostatic regulation would be expected to generate only transient signals that would terminate upon return of the system to the homeostatic set-point. In the absence of homeostatic signaling, N-cadherin and other components of the SJ, as N-cadherin is not known to bind directly to actin, will be maintained in various loaded conformations as SJs mature. This would enable the maintenance, or perhaps even the alteration, of mechanosensitive signaling as loads increase with longer assembly. This may be key to the complex signaling that likely underlies likely complex long-term biological processes, such as synaptic maturation and LTP. Here we have established a correlation between NMII-activity and Src activity. Notably, the RhoGTPases are also known to be subject to mechanosensitive regulation and play key roles in AJ assembly as well as synapse development. In particular, the regulation of RhoGTPases during LTP consolidation is particularly complex. Excitatory stimulation, which promotes NMII-driven synaptic forces (Fig. 5), results in coincident activation of the RhoGTPases, Rac1, Cdc42, and RhoA [76]. RhoA initiates LTP, while Rac1 supports later LTP consolidation [77]. Notably, key regulators of RhoGTPases that preferentially affect each stage of synaptic development, including a variety of GEFs and GAPs as well RhoGDI, have been identified [78]. A major question for future work will be determine if these regulators are subject to mechanosensitive regulation.

In summary, we have created a novel biosensor for visualizing and studying the forces across N-cadherin. We demonstrated that the sensor functions in multiple systems with very different adhesive structures. Comparison of the regulation of loading in N-cadherin during junctional assembly led to the observation of two forms of adhesion strengthening, one consistent with previously observed homeostatic mechanisms and another where loads increased as adhesion assemble. We suggest that the latter form of adhesion strengthening may enable either prolonged, or distinct forms of, mechanosensitive signaling. N-cadherin is a key regulator of many fundamentally important cellular processes and is expressed in a wide variety of load-bearing and/or force-generating cells. Therefore, the development and validation of this novel sensor will enable a wide variety of studies in mechanobiology.

## Materials and Methods

### Generation of NCadTS and Control Constructs

The pcDNA 3.1 NCadTS expression construct was generated using overlap extension PCR [5] using Phusion Polymerase (New England BioLabs). The forward and reverse primers 5’ CCA CAG TAC CCA GTC CGA TCC GCA GCC ATG GTG AGC AAG GGC GAG GAG ACC ACA ATG GGC 3’, and 5’ AAT GAA GTC CCC AAT ATC CCC AGG GTG TGG CTT GTA CAG CTC GTC CAT GCC GAG AGT GAT 3’, respectively, were used to create a mega primer that could be used to insert TSMod-F at amino acid 826 of murine N-Cadherin. To create a suitable template for the insertion primer, N-Cadherin-EGFP (Adgene plamid #18870) was altered using forward and reverse primers, 5’ G GTG GAA TTC GCC ATG TGC CGG ATA GCG GGA GCG 3’ and 5’ ATG CCG CCA CCA CTG CTG ACT CAT CGC CGG CGA G 3’, respectively. The PCR product was inserted into pcDNA3.1 via 5’EcoRI/3’NotI restriction digestion and a standard ligation protocol (T4 DNA Ligase; New England BioLabs) to create pcDNA3.1-Ncadherin. An overlap extension PCR utilizing the TSMod-F-containing megaprimer and pcDNA3.1-NCadherin was performed to create pcDNA3.1 NCadTS.

The pcDNA3.1 NcadTL expression construct was generated by PCR using Phusion Polymerase (New England BioLabs) with pcDNA3.1-NCadTS as a template. The forward and reverse primers 5’ GGC GGA ATT CGC CAT GTG CCG GAT AGC GGG 3’ and 5’ TCC GCG GCC GCT ACT CAC TTG TAC AGC TCG TCC ATG CCG AGA GTG ATC CC 3’, respectively, were used in conjunction to amplify the portion of NcadTL which exists in NcadTS. The PCR product was inserted into pcDNA3.1 via 5’EcoRI/3’NotI restriction and a standard ligation protocol to create pcDNA 3.1 NCadTL.

The pcDNA3.1 NcadVenus expression construct was generated by PCR using Phusion Polymerase (New England BioLabs) with N-Cadherin-EGFP (Adgene plamid #18870) as a template. The c-terminus fluorophore, mVenus was isolated from pcDNA3.1 mVenus using forward and reverse primers 5’ ATA CCG GTC GCC ACC ATG GTG AGC AAG GGC GAG 3’ and 5’ TGC GGC CGC TTA CTT GTA CAG CTC GTC CAT GC 3’, respectively. The PCR product was inserted into N-Cadherin-EGFP via 5’AgeI/3’NotI restriction and a standard ligation protocol. N-Cadherin-Venus was inserted into pcDNA3.1 via 5’EcoRI/3’NotI restriction and a standard ligation protocol to create pcDNA3.1 NcadVenus.

The NcadTS intermolecular FRET controls, pcDNA3.1 NcadTS-dmTFP1 and pcDNA3.1 NcadTS-dVenus, were generated by PCR using pcDNA3.1-NCadTS as the template. Forward and reverse primers were generated for both constructs to induce a Y72L or Y67L amino acid switch within the respective mTFP1 and mVenus chromophores, respectively. Specifically, the forward primers: 5’ CGC GTT CGC CTT AGG CAA CAG G 3’ and 5’ CCT GGG CTT AGG CCT GCA 3’ and reverse primers: 5’ CTG TTG CCT AAG GCG AAC GC 3’ and 5’ TGC AGG CCT AAG CCC AGG 3’ were used for the creation of NcadTS-dmTFP1 and NcadTS-dVenus, respectively. The PCR products were inserted into pcDNA3.1 via 5’NheI/3’XbaI restriction and a standard ligation protocol to create the intermolecular FRET control constructs.

To create lentiviral expression constructs, NCadTS, NCadTL, and intermolecular FRET controls (NcadTS-dmTFP1 and NcadTS-dVenus) were extracted from their respective pcDNA3.1-based expression plasmids via 5’NruI/3’XbaI restriction and ligated into the pRRL lentiviral expression vector (a kind gift of Kam Leong), which had been digested with 5’EcoRV/3’XbaI using standard approaches.

### Cell Culture, Transient Transfections, and Creation of Stable Cell Lines

MOVAS cells (ATCC) were maintained in high-glucose Dulbecco’s Modified Eagle’s Medium (D6429; Sigma Aldrich) supplemented with 10% fetal bovine serum (FBS) (HyClone), 1% antibiotic-antimycotic solution (Sigma Aldrich), and 0.2 mg/ml G418 (Thermo Fisher Scientific, Waltham, MA). A10 cells (ATCC) were maintained in high-glucose Dulbecco’s Modified Eagle’s Medium (D6429; Sigma Aldrich) supplemented with 10% FBS (HyClone, and 1% antibiotic-antimycotic solution (Sigma Aldrich). Cells were grown at 37 °C in a humidified 5% CO2 atmosphere. Transient transfections were performed using Lipofectamine 2000 (Life Technologies) according to the manufacturer’s instructions.

To create lentiviral particles, psPax2 (Addgene plasmid #12260), and pMD2.G (Addgene plasmid #12259) plasmids were co-transfected into HEK293-T cells using Lipofectamine 2000 (Life Technologies). After 4 h, the transfection mixture was exchanged for full media. After an additional 72 h, media containing viral particles was harvested and stored. One day prior to viral transduction, MOVAS or A10 cells were plated in 6-well dishes at a density of 100,000 cells per dish. Cells were transduced with 500 μL viral mixture in full media supplemented with 2 μg/mL Polybrene (Sigma Aldrich) to enhance viral uptake. After three passages, transduced cells were sorted into several groups based on intensity of the fluorescent signal due to expression of the construct. Western blot analysis and immunofluorescent staining were then used to select the population with the optimal expression level.

Low-density hippocampal cultures were prepared from E19 rat embryos as described previously (Zhang *et al.*, 2003) (ScienCell Research Laboratories,), and experiments were carried out in compliance with the Guide for the Care and Use of Laboratory Animals of the National Institutes of Health and approved by the University of Virginia Animal Care and Use Committee (Protocol Number: 2884). Neurons were plated on glass coverslips coated with 1 mg/mL poly-L-lysine at an approximate density of 70 cells/mm^2^ and were transfected with Lipofectamine 2000 (Life Technologies). All neurons used in this study were between days *in vitro* (DIV) 13-22 to assess tension at synapses. Glycine-mediated chemical LTP was performed according to http://science.sciencemag.org/content/305/5692/1972

### Antibodies and Reagents

A mouse monoclonal antibody against gephyrin (Synaptic Systems, clone mAb7, cat. no. 147 011) and a guinea pig polyclonal against VGlut-1 (Synaptic Systems, cat. no. 135-304) were used at concentrations of 1:500 and 1:10000 respectively for immunofluorescence. A mouse monoclonal against PSD-95 (Santa Cruz Biotechnologies, cat. no. sc-32291) was used at a concentration of 1:50-1:100. A rabbit polyclonal antibody against phosphorylated NMDAR Tyr1252 (Abgent, cat. no. AN1077) was used at 1:400. A mouse monoclonal antibody against N-Cadherin (BD Biosciences, cat. no. 610920) was used at a concentration of 1:500. A rabbit monoclonal antibody against β-Catenin (Cell Signaling, cat. no. 8480) was used at a concentration of 1:100. Alexa-conjugated secondary antibodies were from Invitrogen. Blebbistatin (Calbiochem, cat. no. 203391, and Sigma-Aldrich, cat. no. B0560) and used at a concentration of 50 μM for neurons and 25 μM for VSMCs, respectively. Y-27632 (EMD Millipore, cat. no. SCM075) was used at a concentration of 100 μM. Tetrodotoxin and strychnine were purchased from Sigma-Aldrich and reconstituted in dH_2_O. The mGFP-SH2 construct was a gift from Benjamin Geiger [58].

### Immunocytochemistry

Smooth muscle cells (A10 and MOVAS) were fixed in methanol-free 4% paraformaldehyde (PFA) (Electron Microscopy Sciences, cat. no. 15710) for 10 mins diluted in PBS++ (Thermo Fisher Scientific, cat. no. 14040) and then simultaneously blocked and permeabilized with 2% BSA in PBS++ with 0.1% saponin (Alfa Aesar, cat. no. A18820) for 1hr at room temperature. 2% BSA in PBS++ with 0.1% saponin was used for subsequent blocking and antibody incubation. Cells were subsequently imaged in PBS++.

Neurons were fixed in methanol-free 4% formaldehyde ultra-pure EM grade (Polysciences, Inc., cat. no. 18814-10) + 4% sucrose in PBS for 20 min at room temperature and permeabilized with 0.2% Triton X-100 for 5-10 min at room temperature, followed by blocking and antibody incubations in 5-20% normal goat serum. Coverslips were mounted with Vectashield mounting media (Vector Laboratories, cat. no. H-1000). Alternatively, for visualization of N-Cadherin FRET, samples were fixed and imaged in PBS.

### Western Blot Analysis

Western Blots were performed following standard protocols. Briefly, following cell wash and lysis [10% Glycerol, 2 mM EDTA, 250 mM NaCl, 50mM HEPES, 0.5% NP-40, protease inhibitor cocktail (Sigma)], cell lysates were centrifuged for 10 mins at 13000 RPM at 4 °C. 2x Laemmli sample buffer (Bio-Rad Laboratories), was added to supernatants and samples were boiled at 100 °C for 5 mins. Samples were added to mini-protean tgx precast gels (Bio-Rad Laboratories), transferred to PVDF membranes (Bio-Rad Laboratories), and exposed to an N-Cadherin antibody (1:3000) (cat. no. 610920) purchased from BD Biosciences (San Jose, CA). Blots were developed using Supersignal West Pico Chemiluminescent Substrate (Thermo Fisher Scientific), and the signal was detected on X-ray film (Kodak, Rochester).

### Imaging of VSMC Lines

Imaging techniques follow precedent from our previously published work [7]. Samples were imaged at 60x magnification (UPlanSApo 60X/NA1.35 Objective; Olympus) using epifluorescent microscopy on an Olympus inverted fluorescent microscope (Olympus IX83) illuminated by a LambdaLS equipped with a 300W ozone-free xenon bulb (Sutter Instrument). The images were captured using a sCMOS ORCA-Flash4.0 V2 camera (Hamamatsu Photonics). The FRET images were acquired using a custom filter set comprised of an mTFP1 excitation filter (ET450/30x; Chroma Technology Corp.), mTFP1 emission filter (ET485/20m; Chroma Technology Corp), Venus excitation filter (ET514/10x; Chroma Technology Corp), Venus emission filter (FF01-571/72; Semrock) and dichroic mirror (T450/514rpc; Chroma Technology Corp). User-chosen regions of interest (ROI) were photobleached using a 515 nm laser (FRAPPA; Andor Technology) after taking four pre-bleach images. To ensure complete bleaching, 10 laser pulses with a dwell time of 1000 μs per pixel were used. Pre- and post-bleach FRAP images were acquired using the aforementioned Venus excitation and emission filters every 5 seconds until 5 minutes post-bleach. The motorized filter wheels (Lambda 10-3; Sutter Instrument) and automated stage (H117EIX3; Prior Scientific), as well as photobleaching and image acquisition were controlled through MetaMorph Advanced software (Olympus). To maintain a consistent temperature across the sample when conducting live FRAP imaging, an objective heater (Bioptechs) in conjunction with a stage heater (Bioptechs) and heated lid (Bioptechs) were used. Heating equipment was set to 36°C allowed to equilibrate for 20 minutes prior to placing the sample. After the experimental sample was placed, an additional 5 minutes were allowed for the sample temperature to stabilize prior to imaging. During live cell imaging, Medium 199 (Thermo Fisher Scientific) supplemented with 10% FBS (HyClone) and 1% antibiotic-antimycotic solution (Sigma Aldrich) was utilized. Samples were imaged for less than 60 mins to reduce issues due to lack of CO_2_ control.

### FRAP Analysis

Calculation of FRAP recovery was accomplished as previously described [60]. Briefly, user-defined polygons were used to outline a background region outside the cells, the initial position of an unbleached AJs, and the initial position of a bleached AJs. The AJ polygons were refined using the water algorithm [79] with user-optimized parameters, and were automatically moved to account for small (2-10 pixels) movements. For bleached AJs, the position of the polygon was held constant until the intensity of the AJ reached a user-defined threshold, typically 25% of the initial intensity. AJs that grew, shrank, or moved drastically during the experiment were not analyzed.

The recovery curve was normalized to account for initial intensity, background intensity, and global bleaching. Normalizing for global photobleaching was performed by tracking adhesions that were not specifically photobleached, according to previously established methods [80]. The normalized recovery curve was then fit to a single exponential recovery equation:

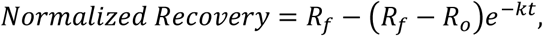

where *R*_*f*_ is the final recovery, *R*_*o*_ is the initial recovery, and *k* is the recovery rate. The half-time of the recovery is determined by *τ*_1/2_ = ln 2/*k*.

### Determination of FRET Efficiency and Molecular Forces in AJs

FRET was detected through measurement of sensitized emission [64] and quantified using custom written code in MATLAB (Mathworks, Natick, MA). A complete description of the FRET efficiency calculations can be found in our previous publications [66, 81]. Prior to FRET calculations, all images were first corrected for uneven illumination, registered, and background-subtracted. Spectral bleed-through coefficients were determined through FRET-imaging of donor-only and acceptor-only samples (i.e. cells expressing a single donor or acceptor fluorescent protein). Donor bleed-through coefficients (*dbt*) were calculated for mTFP1, and acceptor bleed-through coefficients (*abt*) were calculated for mVenus. To correct for spectral bleed-through in experimental data, corrected FRET images (Fc) were generated following the equation:

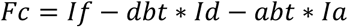

where *If* is the intensity in the FRET-channel, *Id* is the intensity in the donor-channel, and *Ia* is the intensity in the acceptor-channel. After bleed-through correction, FRET efficiency was calculated following the equation:

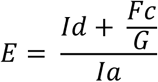

where G is a proportionality constant that describes the increase in acceptor intensity (due to sensitized emission) relative to the decrease in donor intensity (due to quenching) [64]. Towards ensuring that there is a relatively equal stoichiometric abundance of donor and acceptor fluorophores, a donor-per-acceptor (DPA) ratio is calculated and junctions with average DPA ratios less than 0.5 or greater than 1.5 are excluded from analysis.

To quantify characteristics of individual AJs, closed boundaries were drawn by the user based on the unmasked acceptor-channel image around AJs of interest. Junctions for all N-cadherin sensor constructs, imaged both live and fixed in 4% PFA, were segmented based solely on a pixel intensity threshold of 1000 in the acceptor channels. Junctions where remaining pixels summed to less than 2 μm^2^ were excluded from analysis, otherwise average FRET efficiency, average acceptor and donor intensity, AJ area, and DPA were calculated

To measure FRET efficiency in the cell membrane, boundaries were drawn by the user based on the unmasked acceptor-channel image around entire cells except regions with AJs. Pixels with an intensity greater than 500 within the cell boundaries were analyzed for their average FRET efficiency, average acceptor and donor intensity, total area, and DPA. Cell membrane areas junctions with average DPA ratios greater than 0.5 or less than 1.5 were excluded from analysis.

### Imaging of neurons

Confocal images were acquired on a laser scanning Olympus Fluoview 1000 microscope (IX81 base) equipped with a 60X/1.35 NA oil objective (Olympus). For dual-emission ratio imaging of FRET-based sensors, we used the 458 nm line of a multi-Argon ion laser for mTFP1 excitation and SDM510/BA480-495 (mTFP1) and BA535-565 (mVenus/FRET) filters for the collection of mTFP1 and mVenus fluorescence emission, respectively. Green fluorescent probes (GFP) were excited with a 488 nm laser line of a multi-Argon laser, while red probes were excited with the 543 nm laser line of a He-Ne laser; and the far-red probe Alexa647 was excited with the 635 nm line of a diode laser. Fluorescence emission was collected using the following dichroic mirror/filter combinations: SDM560/BA505-525 (GFP), SDM640/BA560-620 (mCherry, RFP, Alexa568 and Rhodamine) and BA655-755 (Alexa647). Fluorescent images were collected in a Z-stack and in sequential line scanning mode using Olympus Fluoview software. Image analysis was performed with Image J software on max intensity Z projections.

### FRET imaging and analysis in Synaptic Junctions

Ratiometric FRET images of max intensity Z projections of the acquired mTFP1 and mVenus images were created using ImageJ for analysis or the Biosensor Processing Software 2.1 available from the Danuser laboratory for removal of background for visualization (University of Texas Southwestern, Dallas, TX) [82]. Analysis of the resulting FRET images was performed with Image J software.

### Statistics

For VSMCs, statistical analyses were performed using JMP Pro 13 software (SAS). Approximately normal data were analyzed using ANOVAs and Student’s t-test to determine if statistically significant differences (p < 0.05) were present between groups. Data sets found to be normally distributed but exhibiting unequal variances as determined by Levene’s test were analyzed with a one-way Welch’s Anova and combined before multiple comparisons were analyzed using the Steel-Dwass test. A summary of relevant non-parametric multiple comparisons for FRET data is available in the supplement (Sup. Table 1). For neurons, statistical analysis was performed using Sigma Plot 13.0. Data was assessed for normality using the Shapiro-Wilk test. Data that had a normal distribution was analyzed using the Student’s t-test to assess statistical significance. Alternatively, the Mann-Whitney Rank Sum Test was used. The statistical test used is indicated in each figure legend.

## Acknowledgements

This work is funded by an Innovative Project Award from the American Heart Association (16IRG27590004).

---

**Supplemental Figure 1.**
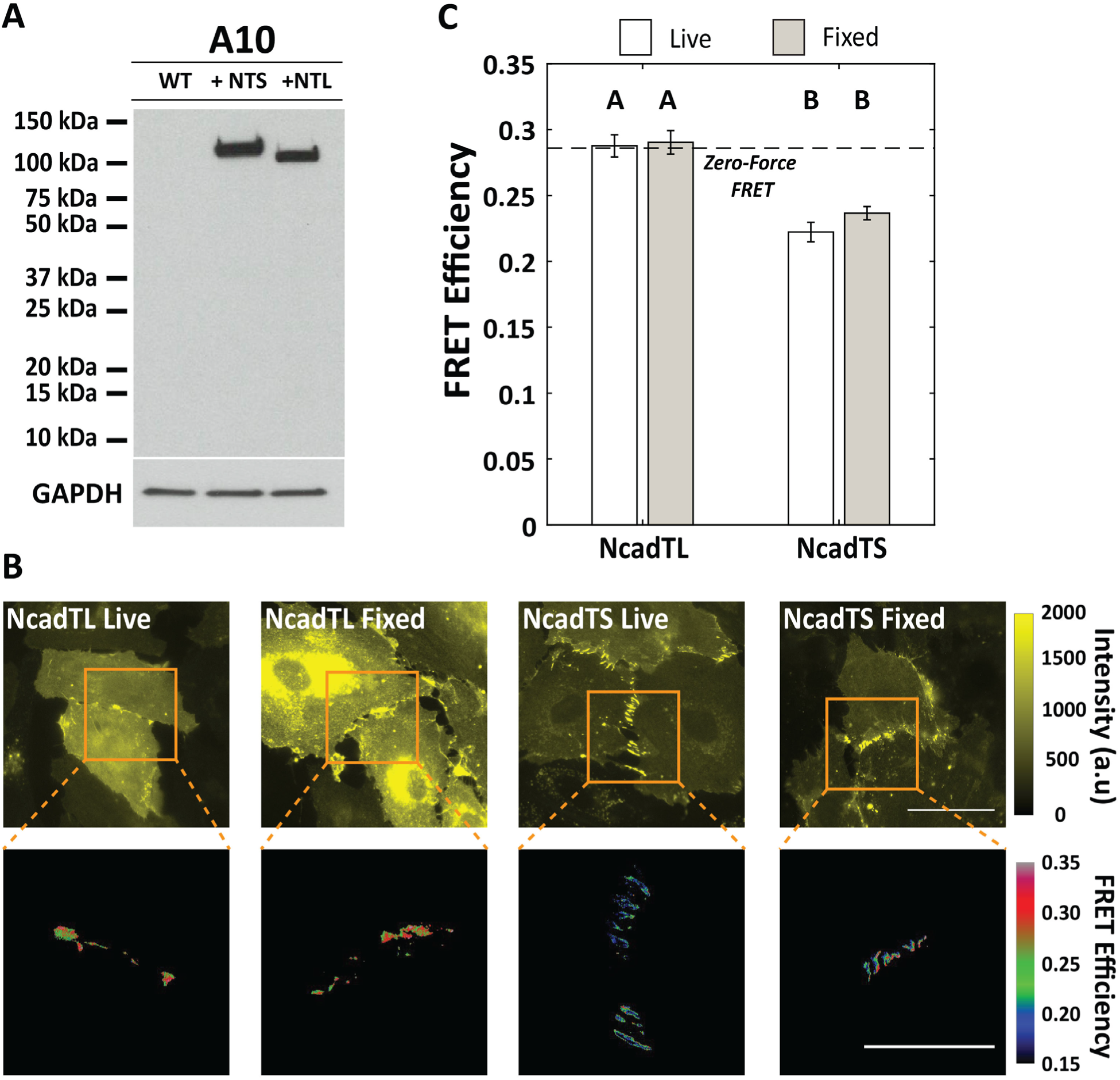
Further validation of N-Cadherin Tension Sensor. **A.** Western Blot of A10 cells which lack endogenous N-Cadherin demonstrate that NcadTS and NcadTL (NTS and NTL, respectively) are produced stably and at the correct size. **B.** A10 cells stably transduced with NcadTL and NcadTS were imaged in live and fixed (4% PFA) conditions and compared to one another. Representative images of NcadTL and NcadTS live and fixed AJs (scale bar = 50μm) and zoom-ins of the FRET readings for the respective, segmented junction (scale bar = 25μm). **C.** Quantification of total data (n = 49, 50, 68, and 31 AJs, respectively). Results from multiple comparisons between groups was conducted using the non-parametric Steel-Dwass test and can be seen in Sup. Table 1. Experimental groups denoted with the same letter are statistically similar, whereas groups with different letters are statistically different.

**Supplemental Figure 2.**
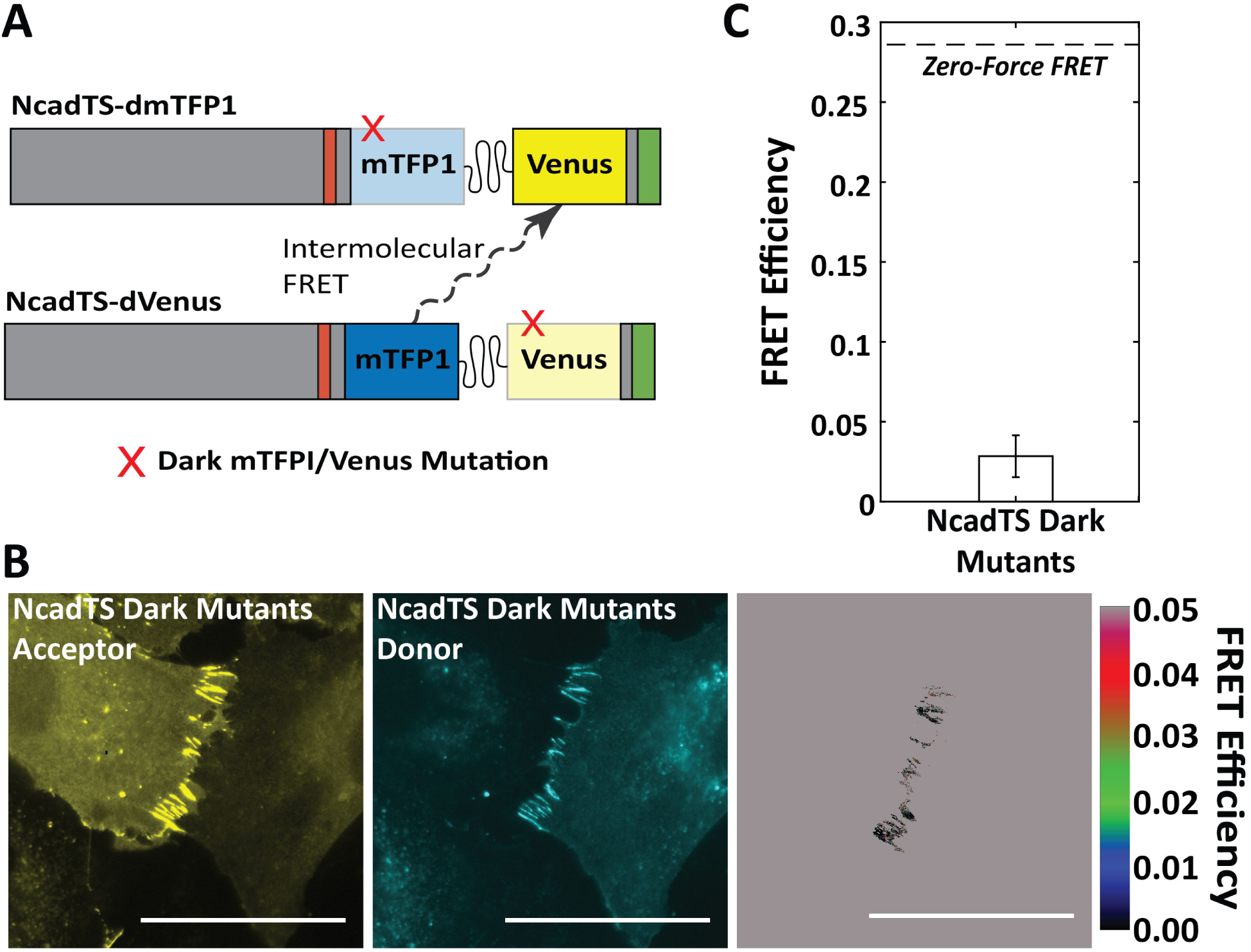
Generation and validation of N-cadherin Intermolecular FRET Sensors. **A.** A dual N-cadherin intermolecular FRET sensor system (NcadTS Dark Mutants) was generated through single amino-acid mutations within the chromophore of mTFP1 and mVenus to produce NcadTS-dmTFP1 and NcadTS-dVenus, respectively. FRET experienced in this system occurs through energy transfer of 2 respective sensors, not within a single sensor. **B.** Representative images of Venus (NcadTS Dark Mutants Acceptor Channel) and mTFP1 (NcadTS Dark Mutants Donor Channel) in the NcadTS-dmTFP1 and NcadTS-dVenus sensors, respectively, as well as the associated FRET at the segmented AJ (Scale bar = 50μm). **C.** Quantification of total data (n = 31 AJs). A10 cells were stably transduced with both dark sensors at equal molarity and FRET was measured and an average FRET reading of ∼3% was measured.

**Supplemental Figure 3.**
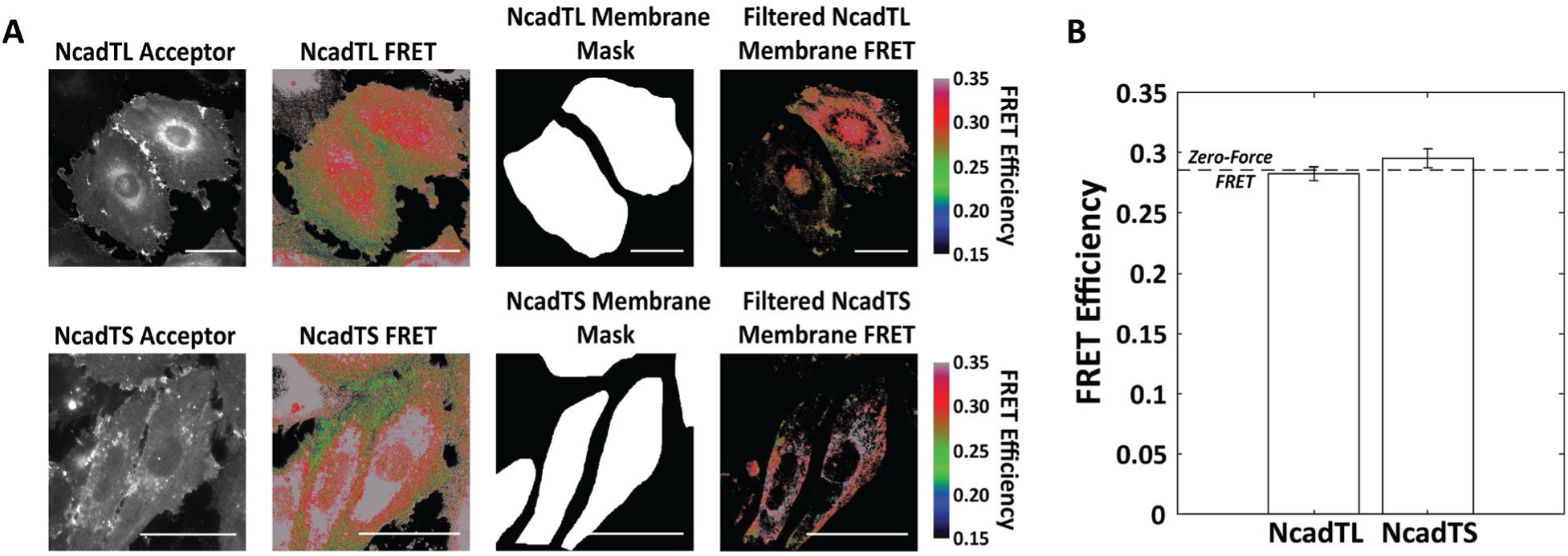
Analysis of Membrane FRET when using the N-Cadherin Tension Sensor. **A.** Process by which membrane FRET was measured. Representative NcadTL and NcadTS acceptor (left) and the calculated FRET efficiencies (center left) are shown. Masks are drawn to isolate entire cells but ignore AJs (center right). Binary masks are applied to the calculated FRET images and pixels that satisfy intensity and DPA thresholds for reliable data are kept (right). (Scale bars = 50μm) **B.** FRET measurements within the cell membrane of A10 cells stably transduced with NcadTL and NcadTS was calculated (n = 102 and 117 regions, respectively). One-sample t-Tests for each sensor against the expected zero-force FRET value (28.6%), reveals no statistical difference for either sensor.

**Supplemental Figure 4.**
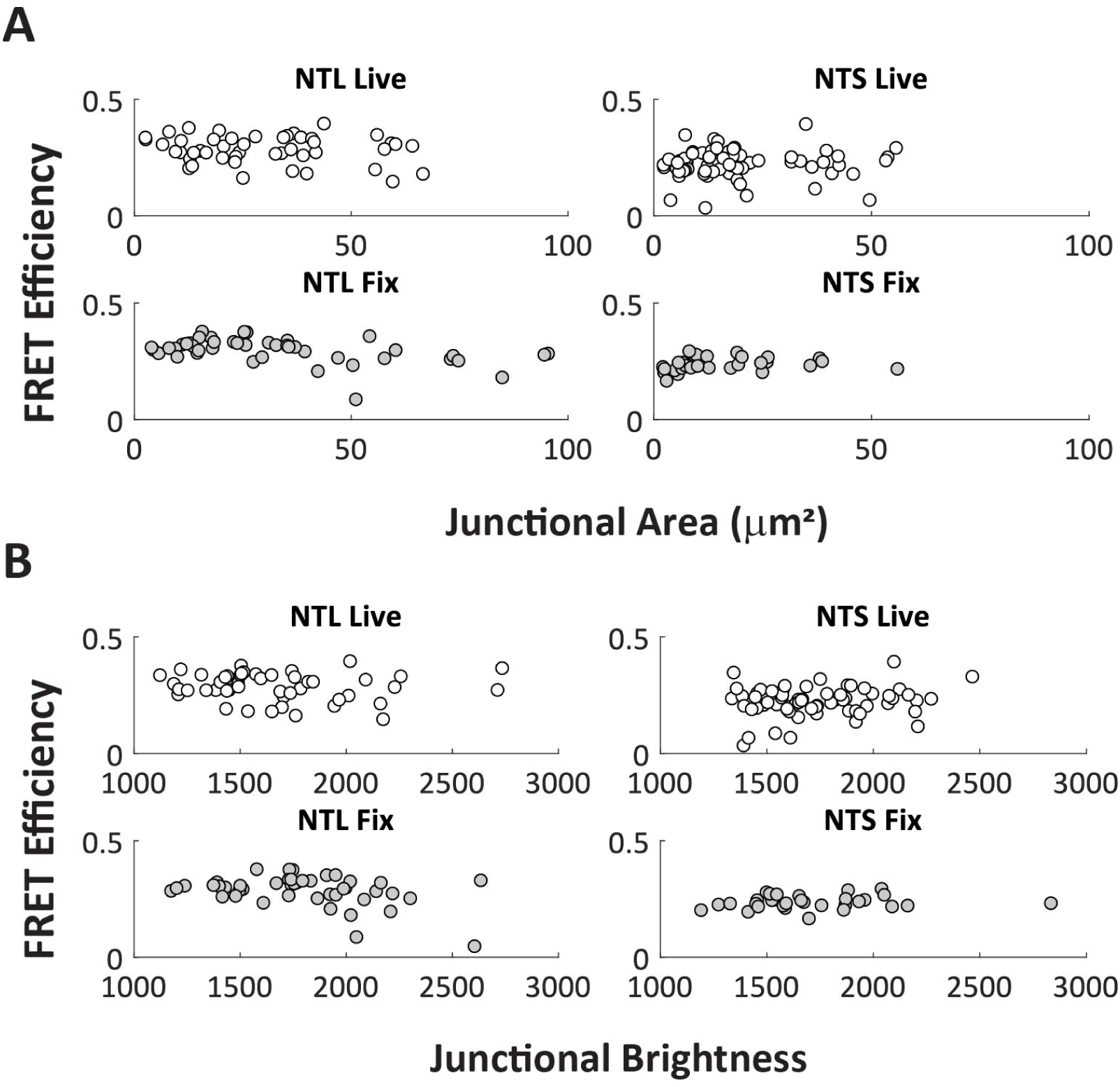
Examining the Relationship between FRET and Junctional Area and Brightness when using the N-Cadherin Tension Sensor. FRET efficiency was analyzed as a function of both (**A.**) junction size (n = 49, 68, 50 and 31 AJs, respectively) and (**B.**) junction brightness (n = 49, 68, 50 and 31 AJs, respectively) when using the NcadTL and NcadTS in both live and fixed imaging paradigms. There does not appear to be a relationship with FRET efficiency based on either size of the junction or the amount of N-cadherin within the junction when conducting either live of fixed imaging.

**Supplemental Table 1.**
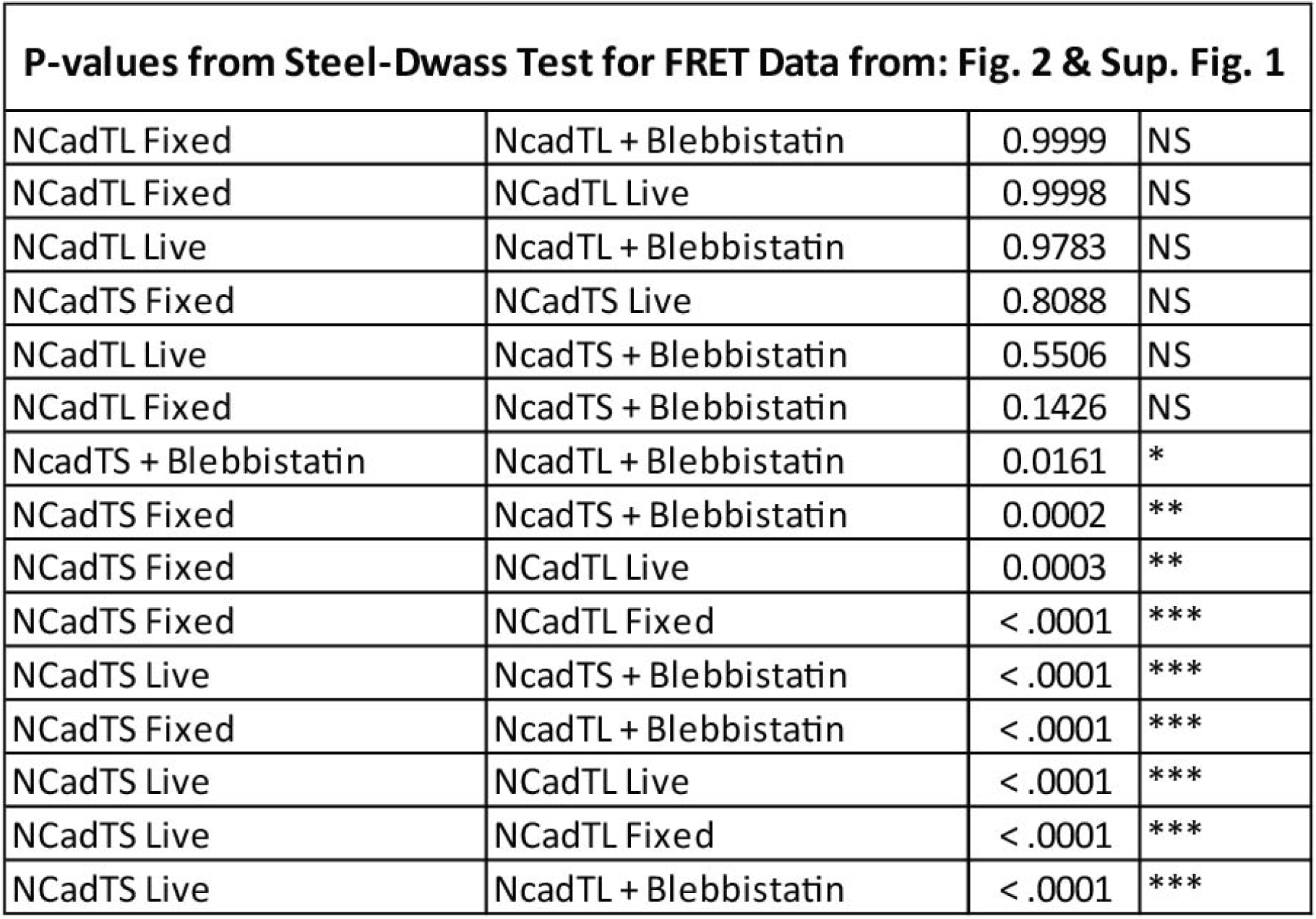
Specific p-values for non-parametric multiple comparisons for data seen in Figure 2 and Supplemental Figure 1.

## Citations

1. Mege, R.M. and N. Ishiyama, Integration of Cadherin Adhesion and Cytoskeleton at Adherens Junctions. Cold Spring Harb Perspect Biol, 2017. 9(5).

2. Seong, E., L. Yuan, and J. Arikkath, Cadherins and catenins in dendrite and synapse morphogenesis. Cell Adh Migr, 2015. 9(3): p. 202–13.

3. Gul, I.S., et al., Evolution and diversity of cadherins and catenins. Exp Cell Res, 2017. 358(1): p. 3–9.

4. Hoffman, B.D. and A.S. Yap, Towards a Dynamic Understanding of Cadherin-Based Mechanobiology. Trends Cell Biol, 2015. 25(12): p. 803–14.

5. Yonemura, S., et al., alpha-Catenin as a tension transducer that induces adherens junction development. Nat Cell Biol, 2010. 12(6): p. 533–42.

6. Leerberg, J.M., et al., Tension-sensitive actin assembly supports contractility at the epithelial zonula adherens. Curr Biol, 2014. 24(15): p. 1689–99.

7. le Duc, Q., et al., Vinculin potentiates E-cadherin mechanosensing and is recruited to actin-anchored sites within adherens junctions in a myosin II-dependent manner. J Cell Biol, 2010. 189(7): p. 1107–15.

8. Borghi, N., et al., E-cadherin is under constitutive actomyosin-generated tension that is increased at cell-cell contacts upon externally applied stretch. Proc Natl Acad Sci U S A, 2012. 109(31): p. 12568–73.

9. Conway, D.E., et al., Fluid shear stress on endothelial cells modulates mechanical tension across VE-cadherin and PECAM-1. Curr Biol, 2013. 23(11): p. 1024–30.

10. Rakshit, S., et al., Ideal, catch, and slip bonds in cadherin adhesion. Proc Natl Acad Sci U S A, 2012. 109(46): p. 18815–20.

11. Conway, D.E. and M.A. Schwartz, Flow-dependent cellular mechanotransduction in atherosclerosis. J Cell Sci, 2013. 126(Pt 22): p. 5101–9.

12. Derycke, L.D. and M.E. Bracke, N-cadherin in the spotlight of cell-cell adhesion, differentiation, embryogenesis, invasion and signalling. Int J Dev Biol, 2004. 48(5-6): p. 463–76.

13. Happe, C.L. and A.J. Engler, Mechanical Forces Reshape Differentiation Cues That Guide Cardiomyogenesis. Circ Res, 2016. 118(2): p. 296–310.

14. Radice, G.L., et al., Developmental defects in mouse embryos lacking N-cadherin. Dev Biol, 1997. 181(1): p. 64–78.

15. Chopra, A., et al., Cardiac myocyte remodeling mediated by N-cadherin-dependent mechanosensing. Am J Physiol Heart Circ Physiol, 2011. 300(4): p. H1252–66.

16. Ganz, A., et al., Traction forces exerted through N-cadherin contacts. Biol Cell, 2006. 98(12): p. 721–30.

17. Lee, E., et al., Deletion of the cytoplasmic domain of N-cadherin reduces, but does not eliminate, traction force-transmission. Biochem Biophys Res Commun, 2016. 478(4): p. 1640–6.

18. Labernadie, A., et al., A mechanically active heterotypic E-cadherin/N-cadherin adhesion enables fibroblasts to drive cancer cell invasion. Nat Cell Biol, 2017. 19(3): p. 224–237.

19. Sun, Z., et al., N-Cadherin, a novel and rapidly remodelling site involved in vasoregulation of small cerebral arteries. J Physiol, 2017. 595(6): p. 1987–2000.

20. Ladoux, B., et al., Strength dependence of cadherin-mediated adhesions. Biophys J, 2010. 98(4): p. 534–42.

21. Chazeau, A., et al., Mechanical coupling between transsynaptic N-cadherin adhesions and actin flow stabilizes dendritic spines. Mol Biol Cell, 2015. 26(5): p. 859–73.

22. Katsamba, P., et al., Linking molecular affinity and cellular specificity in cadherin-mediated adhesion. Proc Natl Acad Sci U S A, 2009. 106(28): p. 11594–9.

23. Vunnam, N., et al., Dimeric states of neural- and epithelial-cadherins are distinguished by the rate of disassembly. Biochemistry, 2011. 50(14): p. 2951–61.

24. Kao, C.Y. and M.E. Carsten, Cellular aspects of smooth muscle function. 1997, Cambridge, U.K.; New York, NY, USA: Cambridge University Press. xv, 293 p.

25. Clark, J.M. and S. Glagov, Transmural organization of the arterial media. The lamellar unit revisited. Arteriosclerosis, 1985. 5(1): p. 19–34.

26. Balint, B., et al., Collectivization of Vascular Smooth Muscle Cells via TGF-beta-Cadherin-11-Dependent Adhesive Switching. Arterioscler Thromb Vasc Biol, 2015. 35(5): p. 1254–64.

27. Gown, A.M., T. Tsukada, and R. Ross, Human atherosclerosis. II. Immunocytochemical analysis of the cellular composition of human atherosclerotic lesions. Am J Pathol, 1986. 125(1): p. 191–207.

28. Frismantiene, A., et al., Cadherins in vascular smooth muscle cell (patho)biology: Quid nos scimus? Cell Signal, 2018. 45: p. 23–42.

29. Sun, Z., et al., N-cadherin, a vascular smooth muscle cell-cell adhesion molecule: function and signaling for vasomotor control. Microcirculation, 2014. 21(3): p. 208–18.

30. Alimperti, S., et al., Three-dimensional biomimetic vascular model reveals a RhoA, Rac1, and N-cadherin balance in mural cell-endothelial cell-regulated barrier function. Proc Natl Acad Sci U S A, 2017. 114(33): p. 8758–8763.

31. Mui, K.L., et al., N-Cadherin Induction by ECM Stiffness and FAK Overrides the Spreading Requirement for Proliferation of Vascular Smooth Muscle Cells. Cell Rep, 2015.

32. Jackson, T.Y., et al., N-cadherin and integrin blockade inhibit arteriolar myogenic reactivity but not pressure-induced increases in intracellular Ca. Front Physiol, 2010. 1: p. 165.

33. Ryu, J., et al., A critical role for myosin IIb in dendritic spine morphology and synaptic function. Neuron, 2006. 49(2): p. 175–82.

34. Lynch, G., C.S. Rex, and C.M. Gall, LTP consolidation: substrates, explanatory power, and functional significance. Neuropharmacology, 2007. 52(1): p. 12–23.

35. Kilinc, D., The Emerging Role of Mechanics in Synapse Formation and Plasticity. Front Cell Neurosci, 2018. 12: p. 483.

36. Hotulainen, P. and C.C. Hoogenraad, Actin in dendritic spines: connecting dynamics to function. J Cell Biol, 2010. 189(4): p. 619–29.

37. Tashiro, A. and R. Yuste, Regulation of dendritic spine motility and stability by Rac1 and Rho kinase: evidence for two forms of spine motility. Mol Cell Neurosci, 2004. 26(3): p. 429–40.

38. Alvarez, V.A. and B.L. Sabatini, Anatomical and physiological plasticity of dendritic spines. Annu Rev Neurosci, 2007. 30: p. 79–97.

39. Yuste, R. and T. Bonhoeffer, Morphological changes in dendritic spines associated with long-term synaptic plasticity. Annu Rev Neurosci, 2001. 24: p. 1071–89.

40. Penzes, P., et al., Dendritic spine pathology in neuropsychiatric disorders. Nat Neurosci, 2011. 14(3): p. 285–93.

41. Uchida, N., et al., The catenin/cadherin adhesion system is localized in synaptic junctions bordering transmitter release zones. J Cell Biol, 1996. 135(3): p. 767–79.

42. Benson, D.L. and H. Tanaka, N-cadherin redistribution during synaptogenesis in hippocampal neurons. J Neurosci, 1998. 18(17): p. 6892–904.

43. Halbleib, J.M. and W.J. Nelson, Cadherins in development: cell adhesion, sorting, and tissue morphogenesis. Genes Dev, 2006. 20(23): p. 3199–214.

44. Fannon, A.M. and D.R. Colman, A model for central synaptic junctional complex formation based on the differential adhesive specificities of the cadherins. Neuron, 1996. 17(3): p. 423–34.

45. Mendez, P., et al., N-cadherin mediates plasticity-induced long-term spine stabilization. J Cell Biol, 2010. 189(3): p. 589–600.

46. Okamura, K., et al., Cadherin activity is required for activity-induced spine remodeling. J Cell Biol, 2004. 167(5): p. 961–72.

47. Nikitczuk, J.S., et al., N-cadherin regulates molecular organization of excitatory and inhibitory synaptic circuits in adult hippocampus in vivo. Hippocampus, 2014. 24(8): p. 943–962.

48. Bian, W.J., et al., Coordinated Spine Pruning and Maturation Mediated by Inter-Spine Competition for Cadherin/Catenin Complexes. Cell, 2015. 162(4): p. 808–22.

49. Togashi, H., et al., Cadherin regulates dendritic spine morphogenesis. Neuron, 2002. 35(1): p. 77–89.

50. Hodges, J.L., et al., Myosin IIb activity and phosphorylation status determines dendritic spine and post-synaptic density morphology. PLoS One, 2011. 6(8): p. e24149.

51. Rex, C.S., et al., Myosin IIb regulates actin dynamics during synaptic plasticity and memory formation. Neuron, 2010. 67(4): p. 603–17.

52. Mysore, S.P., C.Y. Tai, and E.M. Schuman, Effects of N-cadherin disruption on spine morphological dynamics. Front Cell Neurosci, 2007. 1: p. 1.

53. Newell-Litwa, K.A., et al., ROCK1 and 2 differentially regulate actomyosin organization to drive cell and synaptic polarity. J Cell Biol, 2015. 210(2): p. 225–42.

54. Sim, J.Y., et al., Spatial distribution of cell-cell and cell-ECM adhesions regulates force balance while main-taining E-cadherin molecular tension in cell pairs. Mol Biol Cell, 2015. 26(13): p. 2456–65.

55. LaCroix, A.S., et al., Tunable molecular tension sensors reveal extension-based control of vinculin loading. Elife, 2018. 7.

56. Wang, Y., et al., Visualizing the mechanical activation of Src. Nature, 2005. 434(7036): p. 1040–5.

57. Na, S., et al., Rapid signal transduction in living cells is a unique feature of mechanotransduction. Proc Natl Acad Sci U S A, 2008. 105(18): p. 6626–31.

58. Kirchner, J., et al., Live-cell monitoring of tyrosine phosphorylation in focal adhesions following microtubule disruption. J Cell Sci, 2003. 116(Pt 6): p. 975–86.

59. Vicente-Manzanares, M., et al., Non-muscle myosin II takes centre stage in cell adhesion and migration. Nat Rev Mol Cell Biol, 2009. 10(11): p. 778–90.

60. Grashoff, C., et al., Measuring mechanical tension across vinculin reveals regulation of focal adhesion dynamics. Nature, 2010. 466(7303): p. 263–6.

61. Austen, K., et al., Extracellular rigidity sensing by talin isoform-specific mechanical linkages. Nat Cell Biol, 2015. 17(12): p. 1597–606.

62. Sabatini, P.J., et al., Homotypic and endothelial cell adhesions via N-cadherin determine polarity and regulate migration of vascular smooth muscle cells. Circ Res, 2008. 103(4): p. 405–12.

63. Gates, E.M., et al., Improving Quality, Reproducibility, and Usability of FRET-Based Tension Sensors. Cytometry A, 2018.

64. Chen, H., et al., Measurement of FRET efficiency and ratio of donor to acceptor concentration in living cells. Biophys J, 2006. 91(5): p. L39–41.

65. Piston, D.W. and G.J. Kremers, Fluorescent protein FRET: the good, the bad and the ugly. Trends Biochem Sci, 2007. 32(9): p. 407–14.

66. Rothenberg, K.E., et al., Controlling Cell Geometry Affects the Spatial Distribution of Load Across Vinculin. Cellular and Molecular Bioengineering, 2015. 8(3): p. 364–382.

67. Dalva, M.B., A.C. McClelland, and M.S. Kayser, Cell adhesion molecules: signalling functions at the synapse. Nat Rev Neurosci, 2007. 8(3): p. 206–20.

68. Hoffman, B.D., The detection and role of molecular tension in focal adhesion dynamics. Prog Mol Biol Transl Sci, 2014. 126: p. 3–24.

69. Park, M., et al., Recycling endosomes supply AMPA receptors for LTP. Science, 2004. 305(5692): p. 1972–5.

70. Kalia, L.V., J.R. Gingrich, and M.W. Salter, Src in synaptic transmission and plasticity. Oncogene, 2004. 23(48): p. 8007–16.

71. Takasu, M.A., et al., Modulation of NMDA receptor-dependent calcium influx and gene expression through EphB receptors. Science, 2002. 295(5554): p. 491–5.

72. Yap, A.S., K. Duszyc, and V. Viasnoff, Mechanosensing and Mechanotransduction at Cell-Cell Junctions. Cold Spring Harb Perspect Biol, 2018. 10(8).

73. Ringer, P., et al., Multiplexing molecular tension sensors reveals piconewton force gradient across talin-1. Nat Methods, 2017. 14(11): p. 1090–1096.

74. Tzima, E., et al., A mechanosensory complex that mediates the endothelial cell response to fluid shear stress. Nature, 2005. 437(7057): p. 426–31.

75. Bershadsky, A., M. Kozlov, and B. Geiger, Adhesion-mediated mechanosensitivity: a time to experiment, and a time to theorize. Curr Opin Cell Biol, 2006. 18(5): p. 472–81.

76. Hedrick, N.G., et al., Rho GTPase complementation underlies BDNF-dependent homo- and heterosynaptic plasticity. Nature, 2016. 538(7623): p. 104–108.

77. Rex, C.S., et al., Different Rho GTPase-dependent signaling pathways initiate sequential steps in the consolidation of long-term potentiation. J Cell Biol, 2009. 186(1): p. 85–97.

78. Martin-Vilchez, S., et al., RhoGTPase Regulators Orchestrate Distinct Stages of Synaptic Development. PLoS One, 2017. 12(1): p. e0170464.

79. Zamir, E., et al., Molecular diversity of cell-matrix adhesions. J Cell Sci, 1999. 112 (Pt 11): p. 1655–69.

80. Wehrle-Haller, B., Analysis of integrin dynamics by fluorescence recovery after photobleaching. Methods Mol Biol, 2007. 370: p. 173–202.

81. LaCroix, A.S., et al., Construction, imaging, and analysis of FRET-based tension sensors in living cells. Methods Cell Biol, 2015. 125: p. 161–86.

82. Hodgson, L., F. Shen, and K. Hahn, Biosensors for characterizing the dynamics of rho family GTPases in living cells. Curr Protoc Cell Biol, 2010. Chapter 14: p. Unit 14 11 1–26.

